# Allele-specific expression links diverse cis-regulatory mutations to recurrent gene dysregulation in high-risk neuroblastoma

**DOI:** 10.1101/2021.07.08.451695

**Authors:** Arko Sen, Yuchen Huo, Jennifer Elster, Peter E. Zage, Graham McVicker

## Abstract

Neuroblastoma is a pediatric malignancy with a high frequency of metastatic disease at initial diagnosis. Neuroblastoma tumors have few protein-coding mutations but contain extensive somatic copy number alterations (SCNAs) suggesting that mutations that alter gene dosage are important drivers of tumorigenesis. Here we analyze allele-specific expression (ASE) in 96 high-risk neuroblastoma tumors to discover genes with cis-acting mutations that alter dosage. We identify 1,049 genes with recurrent, neuroblastoma-specific ASE, 68% of which lie within common SCNA regions. However, many genes exhibit ASE in copy neutral samples and are enriched for mutations that cause nonsense-mediated decay, indicating that neuroblastoma tumors select for multiple types of mutations that alter gene expression. We also find 24 genes with reduced expression in stage 4 disease that have neuroblastoma-specific ASE that is independent of SCNAs. At least two of these genes have evidence for tumor suppressor activity including the transcription factor *TFAP2B* and the protein tyrosine phosphatase *PTPRH*. In summary, our ASE analysis discovers genes that are recurrently dysregulated by both large SCNAs and other cis-acting mutations in high-risk neuroblastoma.

## Introduction

Neuroblastoma is an extracranial solid tumor of the peripheral sympathetic nervous system which accounts for approximately 8% of all childhood cancers and 15% of childhood cancer mortality^1–6^. Compared to other pediatric malignancies, neuroblastomas harbor few recurrent somatic mutations, and most tumors lack identifiable driver mutations in protein-coding genes at the time of initial diagnosis^7^. Instead, neuroblastoma tumors are characterized by frequent somatic copy number alterations (SCNA). The most common focal SCNA is amplification of the chromosome 2p24 region, including the *MYCN* oncogene, which is associated with high-risk disease and adverse treatment outcomes^8, 9^. Other common SCNAs span tens of megabases and include loss of distal chromosome arms 1p, 3p and 11q and duplication of the distal arm of chromosome 17q^7, 9–12^. These large SCNAs may drive tumorigenesis by altering the expression of multiple tumor suppressors or oncogenes. For example, chromosome 1p deletions affect many potential tumor suppressors including *CHD5*, *CAMTA1*, *KIF1B*, *CASZ1*, *UBE4B*, and *MIR34A*^13–21^. In addition to the common SCNAs described above, neuroblastoma tumors also contain a patchwork of less common SCNAs or Loss of Heterozygosity (LOH) regions. A major challenge in interpreting large SCNAs is that they span dozens of genes, making it difficult to distinguish between driver and passenger genes.

Prior studies of genes with altered dosage in neuroblastoma have largely focused on functional characterization of genes affected by SCNAs while disregarding other dosage-altering mutations. Discovery of important driver genes dysregulated by non-SCNA mutations has been limited because only a small number of whole genome sequences for neuroblastoma are available and it is difficult to determine which noncoding variants affect gene regulation. We hypothesized that genome-wide analysis of allele-specific expression (ASE) could illuminate dysregulated genes in neuroblastoma tumors.

ASE quantifies the difference in expression of two alleles of a gene and can be measured using RNA-seq reads that align to heterozygous sites. Compared to standard differential gene expression analysis, ASE is insensitive to environmental or trans-acting factors, which generally affect both alleles equally. This makes ASE a powerful tool for revealing genes that are affected by cis-acting mutations, including noncoding regulatory mutations that affect sequences such as promoters, enhancers, and insulators as well as protein-coding or splicing mutations that result in nonsense-mediated decay (NMD). Another advantage of ASE is that it is detectable even when the identity of the pathogenic variants causing dysregulation are unknown and it can reveal the effects of rare germline or somatic mutations^22, 23^. In addition to somatic mutations, ASE can also be caused by common germline polymorphisms^24–26^, imprinting^27^ or random monoallelic expression^28, 29^, however these factors are less likely to be involved in tumorigenesis. To differentiate genes which are dysregulated by pathogenic events, the frequency of ASE in disease tissue can be compared to a large panel of normal tissues to identify cancer-specific gene dysregulation^22, 23^. Genome-wide analysis of ASE therefore has the potential to reveal novel tumor suppressor and oncogenes both within and outside SCNAs.

## Results

To discover genes with ASE in neuroblastoma tumors, we obtained exome-seq and RNA-seq data for 96 neuroblastoma tumor samples from the NCI Therapeutically Applicable Research to Generate Effective Treatments (TARGET) project. To estimate ASE in these samples, we implemented a statistical model that utilizes allele-specific read counts at heterozygous sites, while accounting for genotyping errors, sequencing errors and overdispersion of RNA-seq read counts. This model estimates allele imbalance (a_RNA_) for each gene, which is how far the reference allele proportion differs from the expected value of 0.5.

With this method, we identify 8,566 genes with ASE in at least one tumor sample under a false discovery rate (FDR) of 10% (likelihood ratio test). Most genes exhibit ASE in only a single sample (4,664 out of 8,566), however 3,092 genes have ASE in more than one sample, and many genes show highly recurrent ASE in neuroblastoma (948 in 5 or more samples) (Figure 1A, Supplementary Table 1). Since recurrent ASE can result from non-pathogenic factors including common germline polymorphisms^24–26^, imprinting^27^, or random monoallelic expression^28–30^ we compared the frequency of ASE in neuroblastoma to that of normal tissues, obtained from the genotype tissue expression project (GTEx). Specifically, we compared neuroblastoma ASE estimates to those from normal adrenal gland and whole-blood tissues. These tissues were chosen because the adrenal cortex is the tissue of origin for most neuroblastoma tumors and whole-blood has by far the largest number of available samples in GTEx. To illustrate the utility of comparing normal and tumor tissues, we examined ASE for a well-established imprinted gene, *H19*^31^, and for a tumor suppressor gene, *KIF1B*, which is located on chromosome 1p and is frequently deleted in neuroblastoma^16, 32^. As expected, *H19* has very strong ASE in almost all normal and tumor samples (Figure 1B), whereas ASE of *KIF1B* is observed exclusively in neuroblastoma samples (Figure 1C).

**Figure 1:**
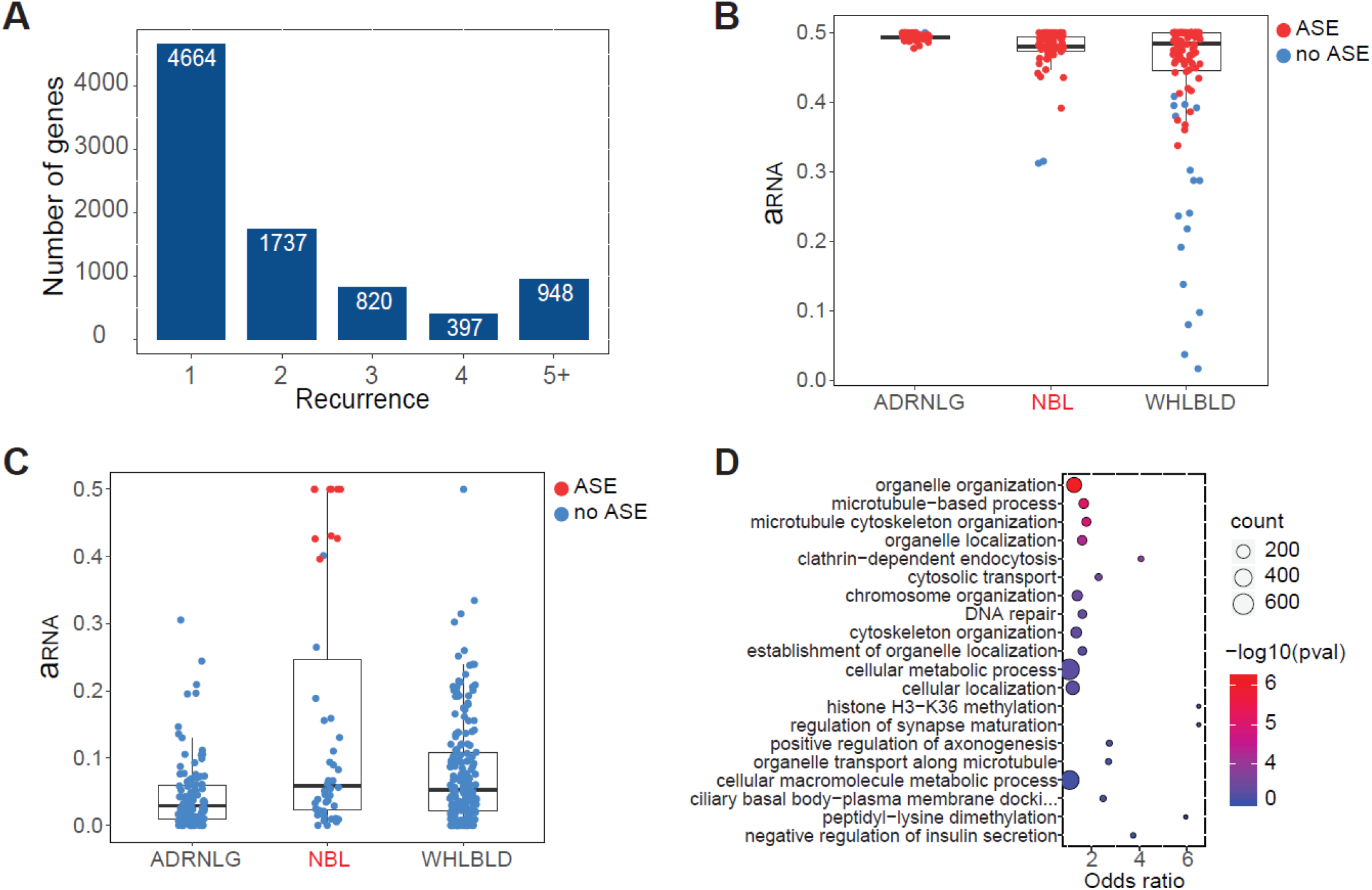
Allele-specific expression analysis in neuroblastoma. **A)** Recurrence of allele-specific expression (ASE) across 96 neuroblastoma samples for genes with ASE in at least one sample (FDR < 0.1). **B)** Estimates of ASE (a_RNA_) for *H19*, an established imprinted gene, in neuroblastoma tumor samples compared to normal adrenal gland and whole blood samples from GTEx. **C)** Estimates of ASE for *KIF1B*, a known neuroblastoma tumor-suppressor gene. **D)** Gene Ontology analysis of 1,049 genes with recurrent ASE in neuroblastoma (NB- ASE genes).

To define a set of genes with neuroblastoma-specific ASE (NB-ASE), we used two filtering criteria: a) genes that are testable for ASE in at least 10 neuroblastoma and 10 adrenal gland or blood samples and b) genes with significant ASE in ≥ 3 neuroblastoma tumors and ≤1 normal tissue (Supplementary Table 2). These criteria resulted in 1,049 NB-ASE genes for downstream analysis. We performed a gene ontology analysis of these genes and found that they are enriched in biological processes frequently dysregulated during tumorigenesis including microtubule-based process (GO:0007017, p-value=6.7e-06), DNA repair (GO:0006281, p-value=3.9e-04), and cellular metabolic process (GO:0044237, p-value=5.1e-04) (Fig 1D, Supplementary Table 3).

SCNAs are a common cause of ASE in tumors^33^, and we hypothesized that many NB-ASE genes would be attributable to large-scale SCNAs that dominate the genetic landscape of neuroblastoma^2, 3, 6, 7^. To determine which NB-ASE genes can be attributed to SCNAs, we adapted our ASE framework to identify SCNAs, which are detectable as large genome segments with allelic imbalance of DNA sequencing reads^34^. While several existing tools leverage read depth to predict SCNAs, these methods have limited precision and report many false positive focal SCNAs^35, 36^. To detect SCNAs, we estimated DNA allelic imbalance from heterozygous sites in windows consisting of 20 consecutive exome capture target regions for tumor (a_tumo*r*_) and normal (a_normal_) samples. We then computed the difference in their absolute values (δ_a_) and performed circular binary segmentation (CBS)^37^ to obtain DNA allelic imbalance for continuous segments which we refer to as the SCNA score (Figure 2A & Supplementary Table 4).

**Figure 2:**
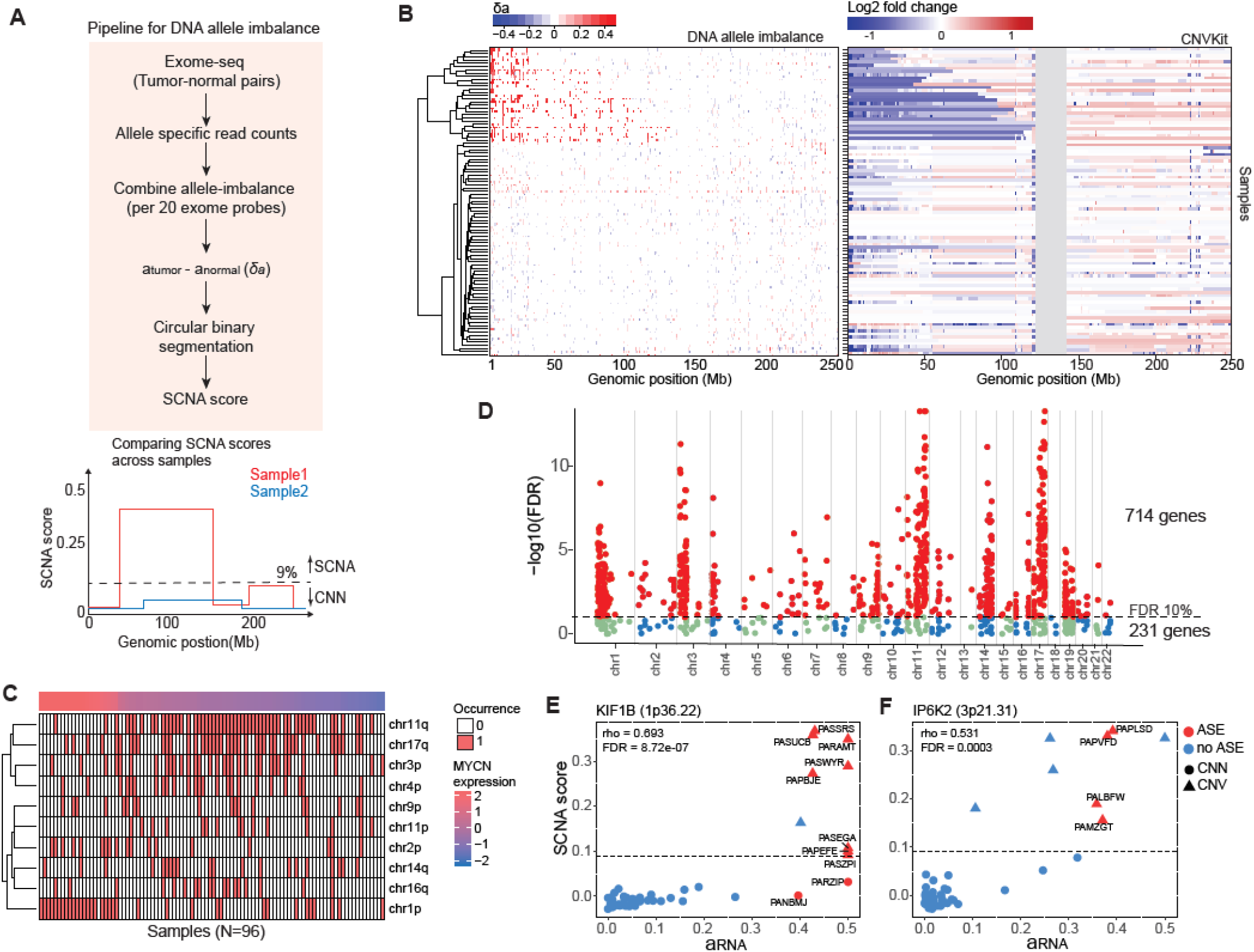
Detecting somatic copy number alterations in neuroblastoma. **A)** Schematic of the DNA allelic imbalance method for detecting somatic copy number alterations (SCNAs). **B)** Difference in DNA allelic imbalance between normal and tumor tissues (*δ*_*a*_) for 96 neuroblastoma patients across chromosome 1 (left panel). Results are compared to log2 fold-change in normalized read coverage between tumor and normal tissues estimated by CNVkit (right panel). **C)** Occurrence matrix of the 10 most frequent SCNAs detected using the DNA-imbalance approach. SCNAs were filtered using SCNA score ≥ 0.09 and annotated based on their cytoband location. Neuroblastoma patients are grouped by *MYCN* gene expression. **D)** Manhattan plot for Spearman’s correlation analysis between ASE (a_RNA_) and SCNA score for 945 NB-ASE genes. **E-F)** Spearman’s rank correlation between allele-specific expression (a_RNA_) and SCNA score for **(E)** *KIF1B*, a tumor suppressor in the chromosome 1p deletion region and **(F)** *IP6K2*, a putative tumor suppressor within the 3p deletion region.

To test our allelic imbalance approach for SCNA discovery we applied it to chromosome 1, which has distal p arm deletions in ∼30% of neuroblastoma tumors^7, 10, 12^. We compared our predictions to CNVkit, which utilizes read depth at exome capture targets to infer copy number and has better sensitivity compared to other methods for SCNA discovery^38^. SCNA breakpoints detected by our method are consistent with those detected by CNVkit, but our predictions are considerably less noisy (Figure 2B). We also compared our results to those from high density single-nucleotide polymorphism (SNP) arrays, which were available for 33 out of the 96 neuroblastoma tumors^39^, and found them to be highly concordant (Supplementary Figure S1). Similar results were obtained for other common SCNAs such as the chromosome 11q deletion region (Supplementary Figure S2).

To further examine the SCNAs in neuroblastoma, we partitioned samples based on *MYCN* expression and found known patterns of SCNA co-occurrence^7^. For example, chromosome 1p and 11q deletions occur most frequently in samples with high and low expression of *MYCN*, respectively (Figure 2C). In addition to the well-characterized SCNAs, we detected less frequent SCNAs across all chromosomes (Figure 2C) including loss of 16q in 18 neuroblastoma tumors (Supplementary Fig S3). This SCNA has not been extensively studied but has been previously reported by comparative genomic hybridization in familial neuroblastomas and some other pediatric cancers such as Wilm’s tumor^40–42^. The 16q deletions are consistently missed by CNVkit (Supplementary Figure S3A), but our deletion predictions for 16q appear to be true positives because they are concordant with both SNP-array predictions and patterns of ASE (Supplementary Figure S3B-E). In combination, these results indicate that DNA allelic imbalance is a powerful approach for the detection of SCNAs in cancer genomes.

To determine whether general patterns of ASE in neuroblastoma can be attributed to SCNAs, we computed Spearman’s correlation between ASE and SCNA score, restricting our analysis to 945 NB-ASE genes that are located within SCNA segments in at least one neuroblastoma sample. Under an FDR of 10%, 68% (714 out of 1,049) of NB-ASE genes are significantly correlated with SCNAs and of these, 58% (411 out 714) are located on the chromosomes with the most frequent SCNAs (chromosomes 1, 3, 11 and 17) (Figure 2D & Supplementary Table 5).

We further examined the relationship between ASE and SCNAs for specific genes in the chromosome 1p and 3p deletion regions that are hypothesized to be tumor suppressors. ASE of *KIF1B* is strongly correlated with SCNA score (Spearman’s rho = 0.69, FDR corrected p-value=9.6e-08), but several samples have strong ASE in the absence of a chromosome 1p deletion (SCNA score ≤ 0.09) (Figure 2E). Thus, in some samples, ASE of *KIF1B* may be caused by other factors such as cis-regulatory mutations, mutations that introduce premature stop codons, or epigenetic changes (Figure 2E). Similar patterns of ASE are present for other putative haplo- insufficient tumor suppressors in the chromosome 1p or 11q deletion regions including *CHD5*^15, 43^, *UBE4B*^13^, *CADM1*^44^ and *ATM*^45^ (Supplementary Figure S4). In contrast, only samples with SCNAs have ASE of *IP6K2*, a putative tumor-suppressor within the 3p deletion region^46, 47^(Figure 2F).

We hypothesized that some ASE events are caused by nonsense-mediated decay (NMD)^48^ introduced by mutations that create premature stop codons including frameshifting indels, nonsense mutations, stop-loss mutations, or mutations that alter splicing. To identify somatic mutations that are likely to cause NMD, we analyzed paired tumor-normal exome-seq data with Variant Effect Predictor (VEP), which collectively labels NMD-causing mutations as ‘high-impact’. We identified 12,122 unique high-impact mutations in the 96 tumor samples, 897 of which are located within 494 NB-ASE genes. Most of these high-impact mutations (85%) are classified as stop gain mutations by VEP (Figure 3A).

**Figure 3:**
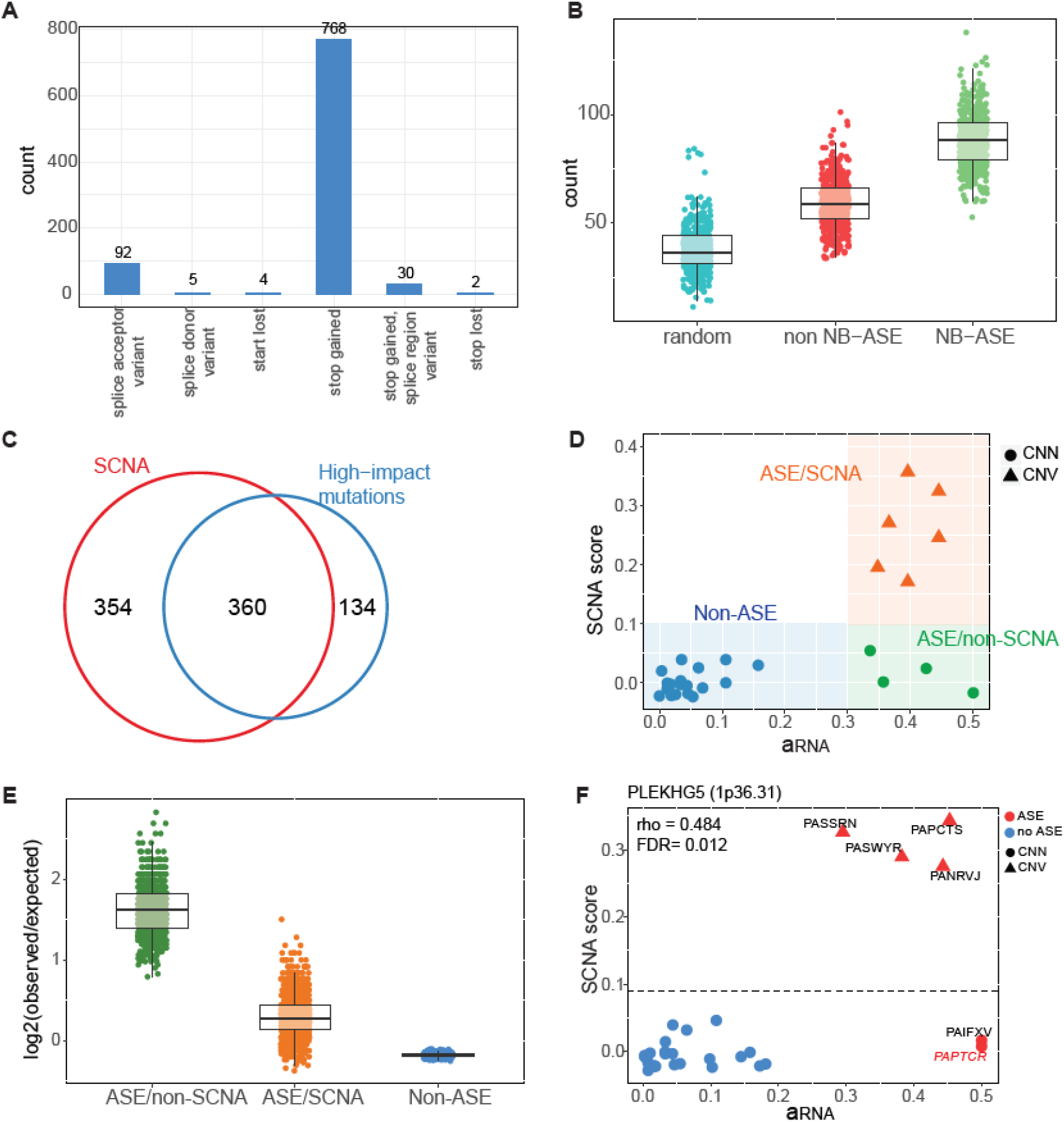
Genes with recurrent allele-specific expression are enriched for high-impact mutations. **A)** The number of variants annotated as high-impact by Variant Effect Predictor (VEP) in genes with recurrent allele-specific expression in neuroblastoma (NB-ASE genes) grouped by functional consequence. **B)** High-impact mutations are enriched within NB-ASE genes. We counted the number of high-impact mutations within 500 samples of 100 genes with samples drawn from (i) a random panel of control genes, (ii) genes with ASE in both neuroblastoma and normal tissues, and (iii) 1,049 NB-ASE genes. **C)** Overlap between genes with correlated ASE and somatic copy number alteration (SCNA) score and NB-ASE genes which contain at least one high-impact somatic mutation. **D)** Graphical representation of three categories of neuroblastoma samples we define for a given gene: samples without ASE (non-ASE); samples with ASE and SCNA (ASE/SCNA); and samples with ASE but no SCNA (ASE/non-SCNA). **E)** The log2 ratio of observed to expected high-impact mutation rate for the three sample categories. Expected rates are estimated by permuting category labels within each gene; points are observed/expected ratios computed across genes after each permutation. **F)** Spearman’s correlation between ASE (a_RNA_) and SCNA score for *PLEKHG5*, a chromosome 1p deletion gene. Two samples have high ASE low SCNA scores. The sample highlighted in red (PAPTCR) carries a premature stop codon mutation.

To determine if NB-ASE genes are enriched for high-impact mutations, we compared the frequency of these mutations in NB-ASE genes to their frequency in two control sets of genes: a) randomly selected genes carrying at least one somatic mutation and b) genes with highly recurrent ASE in both neuroblastoma and normal tissues. We sampled 100 genes, 500 times from each set of genes and counted unique high-impact mutations. The NB-ASE genes contain substantially more high-impact mutations, indicating that NMD is an important driver of neuroblastoma-specific ASE (Figure 3B).

Many genes with correlated ASE and SCNA scores contain high-impact mutations in a subset of samples (Figure 3C), suggesting that NMD is an important mechanism that alters gene dosage in samples that lack SCNAs. To test this hypothesis, we partitioned neuroblastoma samples for each gene into three categories: non-ASE samples; ASE samples with SCNA (ASE/SCNA); and ASE samples without SCNA (ASE/non-SCNA) (Figure 3D). We then calculated the rate of high-impact mutations across all genes and samples in each of the three categories. To generate a null distribution of rates that controls for gene lengths and mutation rate heterogeneity, we permuted the category labels for each gene 1000 times. High-impact mutations occur at a substantially higher rate in ASE/non-SCNA samples compared to other categories (Figure 3E) supporting the hypothesis that in genome regions with frequent SCNAs, gene expression is often altered by NMD or other mechanisms in the samples that lack SCNAs.

An example of a gene which is dysregulated by both NMD and SCNAs is Pleckstrin Homology and RhoGEF Domain Containing G5 (*PLEKHG5*). *PLEKHG5* is located in cytoband 1p36.31 which is frequently deleted in neuroblastoma^49^. Two samples have strong ASE in the absence of SCNAs, one of which contains a C->A heterozygous mutation that introduces a premature stop codon (Figure 3F). The cause of ASE in the other ASE/non-SCNA sample is unknown and could potentially be discovered by analysis of whole-genome sequencing data.

Outside of SCNAs, 104 NB-ASE genes are located in genome regions that are copy neutral across all tumor samples, including *PHOX2B* which is a target of recurrent germline mutations in neuroblastoma^50, 51^. In addition, 231 genes lack significant correlations between ASE and SCNA score even though they overlap SCNAs in one or more samples (Supplementary Table 5). In combination 32% (104 + 231= 335) of NB-ASE genes are not associated with SCNAs, suggesting that other mutational events that alter gene dosage are common in neuroblastoma genomes.

We reasoned that the overall expression of NB-ASE genes could be examined in a larger gene expression dataset where ASE measurements are unavailable. We asked whether the 335 non-SCNA ASE genes are associated with neuroblastoma progression and metastasis, by analyzing the SEQC/MAQC-III Consortium dataset, which contains clinical and microarray expression data for 498 neuroblastoma tumors^52^. Under an FDR of 5% (Student’s t test) and absolute log2 fold-change ≥ 0.5, 35 genes have significantly different gene-expression in stage 4 or metastatic disease compared to other stages (Figure 4). Among them, 11 genes have increased expression and 24 genes have decreased expression in stage 4 disease. Most notably, *MAP7*, *PTPRH, TFAP2B* and *SLC18A1* have more than a 2-fold decrease in expression in stage 4 tumors. We hypothesize that these genes may be important tumor suppressors in neuroblastoma, even though they lie outside of the common SCNA regions of the genome, and we performed further functional analysis of *TFAP2B* and *PTPRH*.

**Figure 4:**
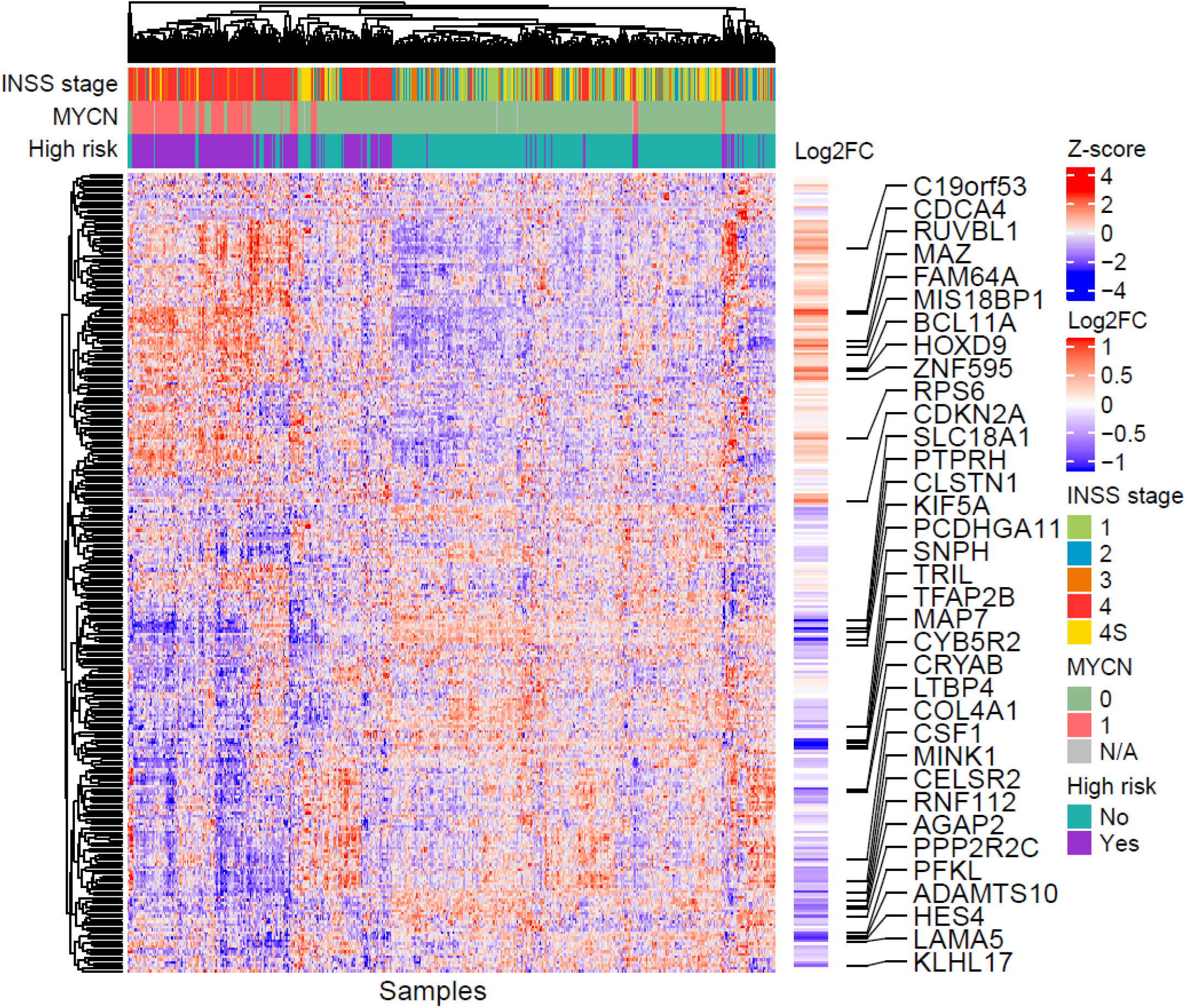
Expression of genes with recurrent allele-specific expression that is not associated with somatic copy number alterations. A hierarchically clustered heatmap of gene expression z-scores from 498 neuroblastoma tumors in the SEQC/MAQC-III Consortium. Of the 335 NB-ASE genes that were not associated with somatic copy number alterations (SCNAs), 294 were covered by at least one expression probe. Samples are labeled by clinical characteristics: *MYCN* amplification status (amplified, non-amplified and unknown); high-risk tumor (yes or no); International Neuroblastoma Staging System (INSS) (1, 2, 3, 4 and 4S). Difference in gene-expression between stage 4 and all other stages is indicated as log2 fold-change (log2FC). The 35 genes with absolute log2FC ≥ 0.5 and FDR corrected p-value ≤ 0.05 from Student’s t test are labelled.

We first investigated *TFAP2B*, which is a retinoic acid-induced transcriptional activator that mediates noradrenergic neuronal differentiation of neuroblastoma cells in vitro^53, 54^. *TFAP2B* has ASE in 3 out of 31 testable neuroblastoma samples, has no evidence of ASE in adrenal gland tissues (0 out of 12 testable samples), is not expressed in whole blood, and is copy neutral in all patient samples (Supplementary Figure S5A). Consistent with the above observations, the samples with ASE of *TFAP2B* have strong allelic imbalance of RNA-seq reads at heterozygous sites, but no allelic imbalance of exome-seq reads, indicating that the ASE is not due to SCNAs (Supplementary Figure S5B). Dysregulation of *TFAP2B* in neuroblastoma cells has previously been associated with aberrant promoter-methylation^54^, so we investigated DNA methylation as a potential mechanism. Using estimates of promoter methylation computed from the Human Methylation 450K array, we found that *TFAP2B* is one of one of only 6 NB-ASE genes with a significant correlation between ASE and promoter-methylation under an FDR of 10% (Spearman’s correlation coefficient = 0.60, FDR-corrected p-value 0.09) (Supplementary figure 6 & Supplementary Table 8). Furthermore, one patient sample (PASNZU) has near-complete methylation (>75%) of the *TFAP2B* promoter, which is associated with loss-of-expression of both alleles (Supplementary Figure S5C-E, Supplementary Table 7-8). In the SEQC/MAQC-III Consortium data, *TFAP2B* expression is decreased in stage 4 or metastatic neuroblastomas (Supplementary Figure S5F) and low expression of *TFAP2B* is associated with worse event-free survival outcomes in non-*MYCN* amplified neuroblastoma patients (Supplementary Figure S5G). Collectively these observations are consistent with earlier findings^54^, and strongly suggest that *TFAP2B* is a tumor-suppressor in neuroblastoma with decreased expression in the presence of promoter methylation.

We next investigated *PTPRH*, (Protein Tyrosine Phosphatase Receptor Type H), which is a member of a large family of receptor tyrosine phosphatases and a critical regulator of apoptosis and cell motility^55, 56^. In neuroblastoma, 5 out of 68 testable samples exhibit ASE, compared to 1 out 139 testable adrenal gland tissues from GTEx (Fisher Exact Test p-value= 0.015) (Figure 5A); *PTPRH* is not expressed in whole blood. *PTPRH* is located on chromosome 19q, which rarely undergoes SCNA in neuroblastoma, and ASE of *PTPRH* is not correlated with SCNA score (Figure 5B). In addition, the RNA-seq reads in ASE samples exhibit strong allelic imbalance but there is no allelic imbalance in exome-seq reads, confirming that ASE of *PTPRH* is not attributable to large or focal SCNAs (Figure 5C). ASE of *PTPRH* is negatively correlated with gene-expression (Spearman’s correlation coefficient = -0.57, FDR corrected p-value = 3.48e-07) indicating that ASE reflects loss of expression of one allele, potentially due to regulatory or other cis-acting mutations (Figure 5D & Supplementary table 7). Gene expression of PTPRH is substantially reduced in stage 4 tumors and reduced expression of this gene is associated with worse event-free survival outcomes (Figure 5E & F). To further test the function of *PTPRH*, we performed shRNA knockdown experiments in the neuroblastoma cell lines SK-N-SH and SK-N-BE(2), which are *MYCN* non-amplified and amplified, respectively. Knockdown of PTPRH increases proliferation in both cell lines and cellular migration in the SK-N-SH cell line. These results support the hypothesis that *PTPRH* is a MYCN-independent tumor suppressor (Figure 6A & B).

**Figure 5:**
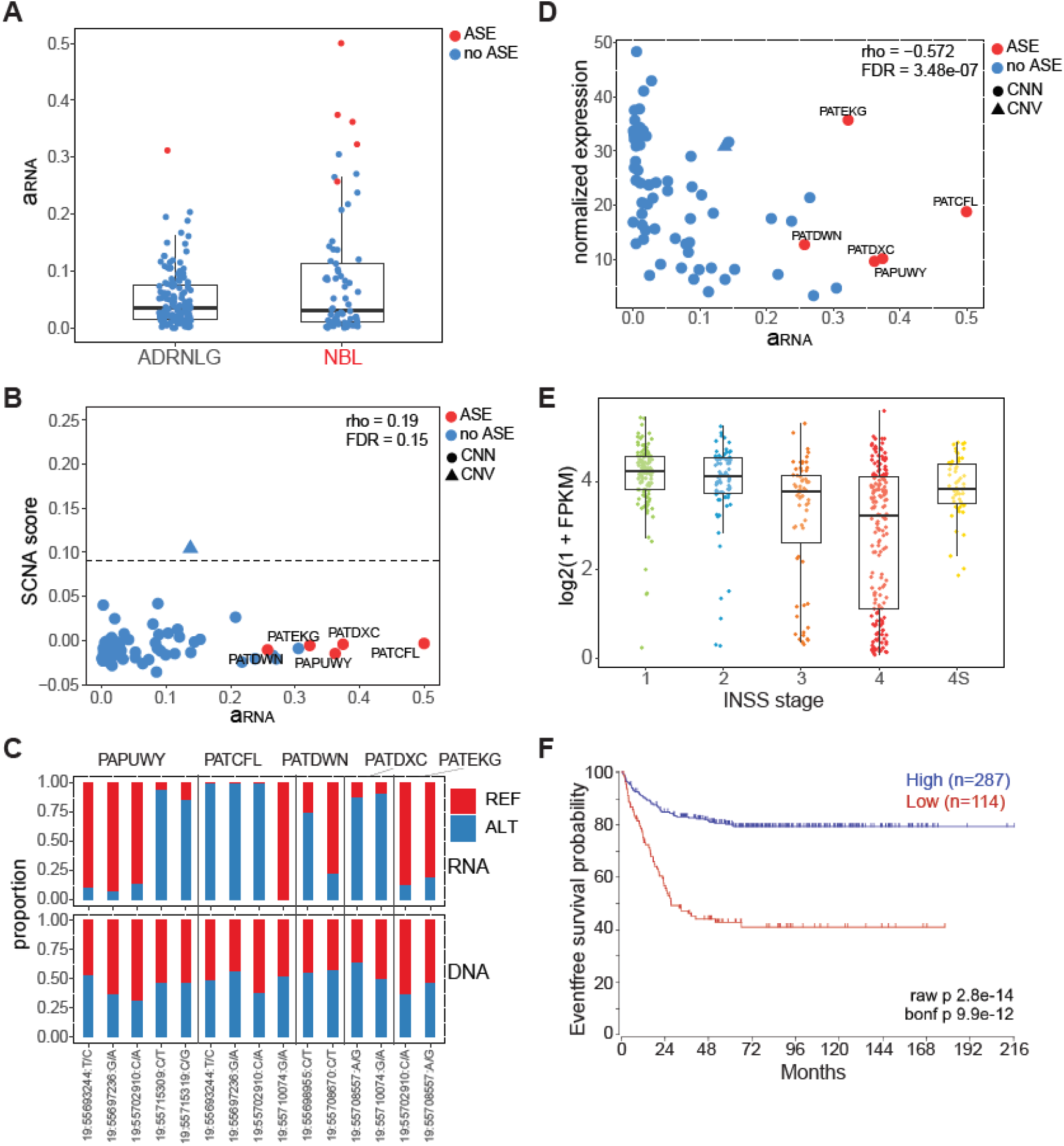
Genomic profiling of *PTPRH*. **A)** ASE (a_RNA_) of *PTPRH* in neuroblastoma and adrenal gland tissues. *PTRPH* has detectable ASE in 5 out of 68 neuroblastoma patients and 1 out of 139 Adrenal gland tissues (Fisher Exact test p-value = 0.015). **B)** Spearman’s correlation between ASE (a_RNA_) for *PTPRH* and SCNA score for the overlapping genomic segments. **C)** Reference and alternate allele proportions for RNA-seq and exome-seq reads at heterozygous sites within the *PTPRH* gene in 5 samples with significant ASE. **D)** Spearman’s correlation between ASE (a_RNA_) and gene expression for *PTPRH.* **E)** Normalized gene expression of *PTPRH* across different disease stages for 498 neuroblastoma patients from the SEQC/MAQC-III Consortium data. **F)** Kaplan Meier survival analysis for MYCN non-amplified patients from the SEQC/MAQC-III Consortium data set.

**Figure 6:**
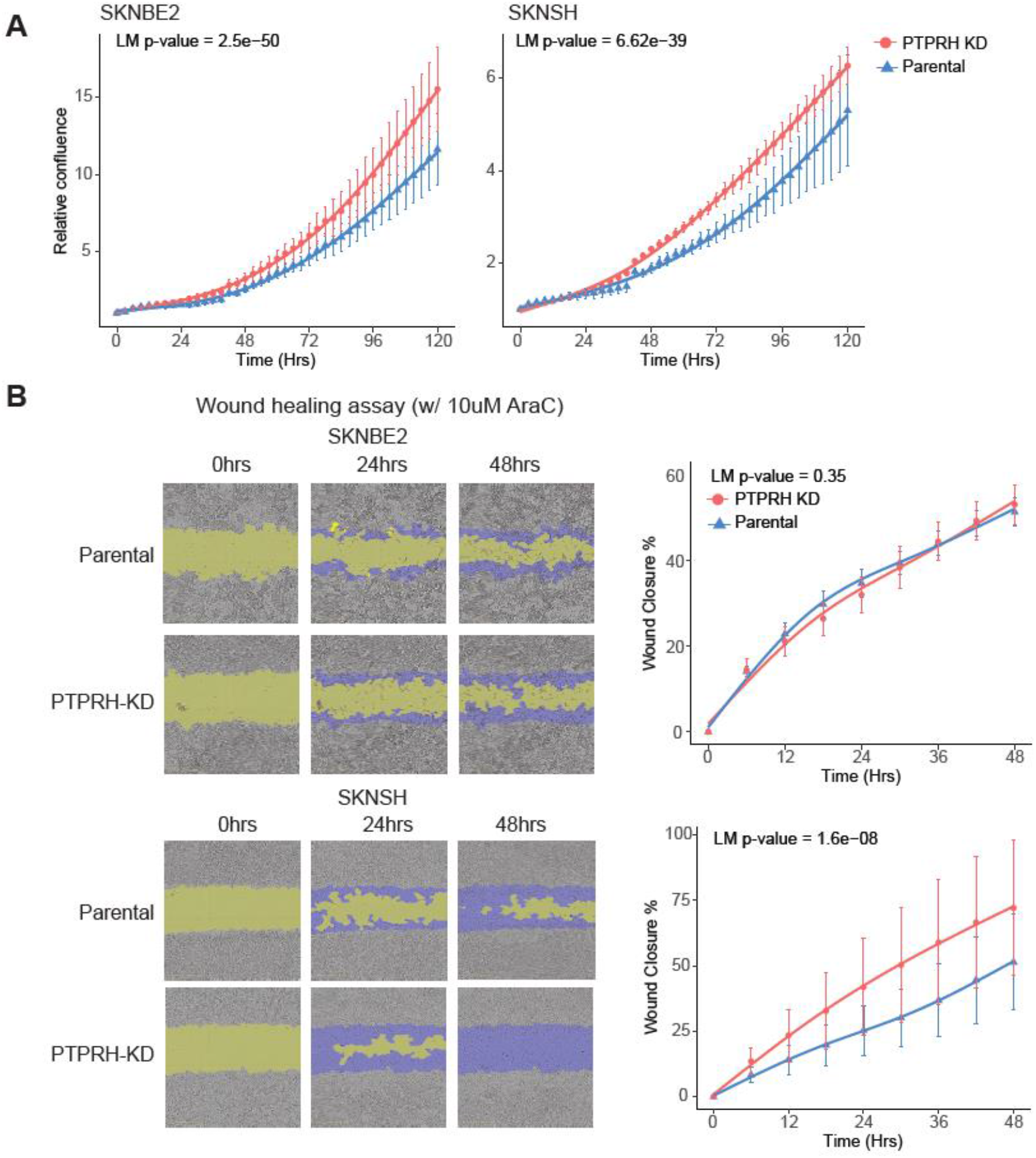
PTPRH knockdown in neuroblastoma cell lines increases cellular proliferation and migration. **A)** Change in relative confluence (i.e., confluence at each time point normalized to confluence at time 0) for parental and PTPRH knockdowns (KDs) in SK-N-BE(2) and SK-N-SH cells. 6 replicates for each condition were used for the SK-N-BE(2) cell line and SK-N-SH cell line. **B)** Left panels: Representative images for scratch wound assays used to measure cellular migration for parental and PTPRH-KD in SK-N-BE(2) and SK- N-SH cells. Right panels: Wound closure percentage for parental and PTPRH-KD in SK-N-BE(2) and SK-N-SH cells over 48 hours. Each time point had 9 replicates. In both **A** and **B** error bars indicate +/- one standard deviation around the mean across replicates at each time point. The lines are fits from cubic spline regression using 3 knots, and the P-value is from an F-test for a difference in splines between parental and KD cells.

## Discussion

Our study leveraged allelic imbalance of RNA and DNA sequencing reads to discover genes with recurrent ASE and delineate SCNA regions in neuroblastoma genomes. Neuroblastoma genomes contain a surprisingly large number of genes with recurrent ASE; however, the majority of ASE events can be attributed to SCNAs that span tens of megabases. Our ASE analysis reveals that, in some samples, genes within recurrent SCNA regions are dysregulated by non-SCNA events including mutations that cause nonsense-mediated decay. Non- SCNA ASE events are present in putative tumor suppressors previously studied in the context of recurrent SCNAs including *KIF1B, PLEKHG5*, *UBE4B*, *CHD5*, *CADM1*, and *ATM*. With larger sample sizes, ASE could potentially be utilized to distinguish passenger genes from driver genes within recurrent SCNA regions.

Discovery of genes by ASE is distinct from standard differential gene expression analysis because it specifically identifies genes impacted by cis-acting events, even when the exact events (e.g. mutations) are not known. In contrast, most genes discovered by differential gene expression may be downstream of the causal mutational events or have altered expression due to environmental differences. Outside of recurrent SCNA regions, we discovered 335 genes that are recurrently dysregulated in neuroblastoma. These genes include *TFAP2B*, *MAP7*, *PTPRH* and *SLC18A1*, which have substantially lower expression in stage 4 disease.

*TFAP2B* is important for noradrenergic neuronal differentiation of neuroblastoma cells in vitro and is dysregulated by aberrant promoter methylation^54^. Our independent validation of this finding is additional evidence that *TFAP2B* is an important tumor suppressor in neuroblastoma. *PTPRH* belongs to a group of receptor tyrosine phosphotases which reduce phosphorylation of Akt and its cellular substrates such GSK-3α, or GSK-3β^56^. *PTPRH* may inactivate Akt and promote apoptosis in cancer cells. In addition, overexpression of *PTPRH* has been demonstrated to disrupt actin-based cytoskeleton as well as inhibit cellular responses promoted by integrin-mediated cell adhesion, including cell spreading on fibronectin, growth factor-induced activation of extracellular signal-regulated kinase 2, and colony formation^55^. Thus, prior studies and our discovery that *PTPRH* exhibits recurrent ASE in neuroblastoma suggest that it functions as a tumor suppressor.

This study provides a framework for studying the impact of non-coding mutations in neuroblastoma and provides evidence that multiple mutational processes work in concert to dysregulate gene expression and drive neuroblastoma tumorigenesis. Future efforts focused on whole genome sequencing and noncoding variants in accessible chromatin regions may help uncover novel regulatory variants and further illuminate the genetic landscape of neuroblastoma.

## Methods

### Datasets

Next Generation Sequencing (NGS) data for neuroblastoma patients were obtained from the Therapeutically Applicable Research to Generate Effective Treatments (TARGET) initiative^7^. Our dataset consisted of RNA sequencing for 143 tumors and paired tumor-normal exome sequencing for 97 neuroblastoma patients. Out of the 97 samples with both RNA-seq and exome-seq data, 87 also had Illumina Infinium Human Methylation 450K data and 33 had HumanHap 550K BeadChIP (SNP-array) data. We also obtained 175 adrenal gland and 369 whole blood samples from the GTEx Consortium and used them as a normal ASE reference set^57^.

### Quality control

To ensure that NGS data from the same patient are properly paired, we compared the RNA-seq and exome-seq data from 97 neuroblastoma tumors using NGSCheckMate^58^. Based on this analysis, we found that 1 sample had mismatched RNA-seq and exome-seq data and we removed this sample from the study. Our final dataset for ASE analysis consisted of RNA-seq and exome-seq data from 96 neuroblastoma patients.

### Variant calling pipeline

We aligned exome-seq reads to the reference genome (hg19) using BWA-MEM with default parameters^59^. Then we generated GVCF files for each sample using the GATK *HaplotypeCaller* (4.1.1) and performed joint genotyping using GATK *GenotypeGVCFs*. We extracted single nucleotide polymorphism (SNPs) using GATK *SelectVariants* command and recalibrated variant quality scores with GATK variant quality score recalibration (VQSR) pipeline. The filtered and processed SNPs were used for downstream analyses.

### Somatic mutation discovery pipeline

We used Mutect2 from GATK (4.1.1) to compare the mutation profile from exome-seq data for 96 neuroblastoma tumor and normal whole blood samples^60^. We filtered somatic mutations from Mutect2 using the GATK recommended filtering pipeline (https://gatk.broadinstitute.org/hc/en-us/articles/360035531132). To determine the functional consequence of somatic mutations and to assign mutations to respective genes we further analyzed individual somatic mutations in each sample using variant effect predictor^61^.

### Read depth-based detection of somatic copy number alternations

SCNAs in neuroblastoma were detected using our DNA allelic imbalance method described below, and a read depth-based method, CNVkit^38^. Briefly, CNVkit uses exome-seq reads to calculate log2 copy ratios across the genome for tumor-normal pairs. SCNAs for large chromosomal regions are then detected by combining log2 copy ratios across adjacent genomic regions using Circular Binary Segmentation (CBS). For this study, we processed aligned exome-seq reads with the *batch* option from CNVkit using the *--drop-low-coverage* parameter to control for low coverage exome targets. Heatmaps showing CNV calls for all samples were generated using CNVkit’s *heatmap* script.

### RNA-seq alignment and processing

We aligned RNA-seq reads end-to-end to the reference genome (hg19) using STAR (2.5.3a)^62^. The aligned reads were filtered to those with mapping quality ≥ 20 using samtools (1.9)^63^. Reads mapping to each gene were counted using featureCounts (1.6.3) for GENCODE (v28) genes^64^. Gene counts were converted to Fragments Per Kilobase Per Million (FPKM) using DESeq2 (1.22.2)^65^. Finally, the FPKM matrix was quantile normalized using the preprocessCore (1.44) package and z-score transformed.

### Estimating DNA allelic imbalance using exome-seq

We realigned exome-seq reads from tumor and normal samples to the reference genome (hg19) using BWA-ALN and corrected the sequencing reads mapping to heterozygous sites for mapping bias using WASP^66^.

Then, we obtained allele-specific read counts at heterozygous sites for normal samples using the *CollectAllelicCounts* function from GATK (4.1.1). We assumed that most heterozygous sites in the normal sample are germline polymorphisms and obtained allele-specific read counts at the same genomic positions for the matched tumor samples. Using the same positions facilitates direct comparison of allele proportions between tumor and normal samples.

To model DNA allelic imbalance over large genomic segments, we sorted exome capture target regions by their genomic coordinates and grouped 20 consecutive target regions into genomic bins. Next, we assigned the heterozygous sites which were covered by at least 10 reads to the genomic bins. To ensure robust regional DNA allele imbalance estimates, we retained genomic bins with ≥10 overlapping heterozygous sites for analysis. We assumed that the reference allele count at heterozygous sites is beta-binomially distributed. The likelihood at heterozygous site *i* within a genomic bin is then defined by:

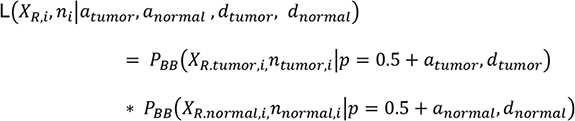

Where X_R,i_ is the observed reference allele count at site *i*, and n_i_ is the total count of reads matching the reference or alternate allele (n_i_=X_R,i_+X_A,i_) at site i, *a* is the allelic imbalance and is defined over the range [- 0.5,0.5], and *d* is the dispersion parameter.

We estimate the dispersion parameter by fixing *a* for tumor and normal samples to 0 and setting *d* to the value that maximizes the total likelihood across all heterozygous sites. We estimate separate values of *d* for tumor and normal samples.

A given bin contains multiple heterozygous sites and we do not know the phasing of the alleles. To account for the uncertainty in phase, we assume that the reference alleles at heterozygous sites within a bin have a 50% probability of being on the same haplotype as the reference allele at the first heterozygous site within the bin. The combined likelihood of the *m* sites within a bin is then:

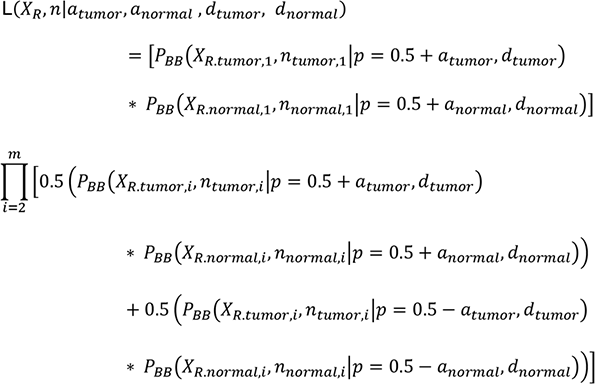

We used maximum likelihood to estimate *a* for tumor and normal samples per *bin* and calculate the difference (δ_a_) as:

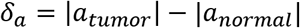

Finally, to create contiguous segments of allelic imbalance, we performed Circular Binary Segmentation (CBS) on the estimates of δ_a_ using DNAcopy (1.56.0)^37^. We call the aggregate value for δ obtained from CBS the SCNA score. Plots for δ_a_ and SCNA scores were generated using Gviz (1.26.5), ComplexHeatmap (1.20.0) and gplots (3.0.1.1)^67^.

### Estimating allele-specific expression per gene using RNA-seq reads

The mapped high-quality reads from RNA-seq alignments were corrected for mapping bias using WASP. Allele-specific read counts at heterozygous positions were obtained using WASP’s *bam2h5.py* script^66^. Only heterozygous positions which pass Variant Quality Score Recalibration (*VQSR*) from GATK (4.1.1) were used for downstream analysis.

Genotyping errors and sequencing errors can create a false signal of allelic imbalance. To control for genotyping errors, we calculate the error rate directly from the genotype quality (GQ) scores from GATK:

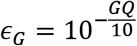

Bi-allelic sites within an individual can generate reads mapping to reference or alternate alleles. However, some sites show a small fraction of reads that match neither the reference nor alternate allele which we refer to as “other” reads or *X*_*O*_. We approximate the sequencing error rate, ɛ_S_, using the *X*_*O*_ read counts. To account for the fact that 1/3 of sequencing errors will be counted as alternate or reference alleles, we scale the error rate estimate, by 3/2:

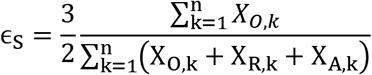

When a genotyping error occurs, a heterozygous site is equally likely to be homozygous reference (0/0) or alternate (1/1). Therefore, conditional upon a genotyping error having occurred, the likelihood at site *i* is defined as:

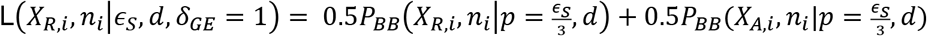

and conditional on no genotyping error the likelihood is:

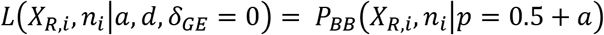

where P_BB_ is the beta binomial probability distribution function, *n*_*i*_ is the total count of reads matching reference or alternative (*n*_*i*_ = *X*_*R*,*i*_ + *X*_*A*,*i*_) at heterozygous site *i*, *a* is the allelic imbalance parameter defined over the range [-0.5,0.5], *d* is the dispersion parameter and *δ_GE_* is an indicator variable that is 1 when a genotyping error has occurred and 0 otherwise. To find a maximum likelihood estimate of *d*, we set *a* to 0 and optimize *d* over all heterozygous sites overlapping exons. Finally, a single gene might contain multiple heterozygous sites which need to be combined to estimate ASE for a gene. We define the likelihood of the read counts for the first heterozygous site within a gene as:

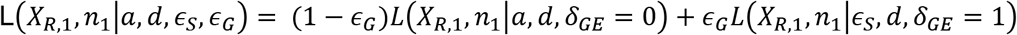

For subsequent heterozygous sites in the same gene, we do not know the phase of the alleles with respect to the first heterozygous site. We assume that the reference and alternative alleles are equally likely to be on the same haplotype as the reference allele at the first site. The combined likelihood of all sites within a gene is then defined as:

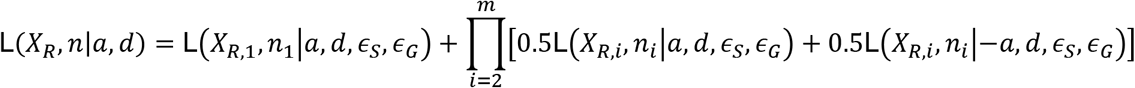

We obtain a maximum likelihood estimate of *a* for each gene under the alternative model of allelic imbalance. Then we use a likelihood ratio test to compare the alternative model to the null model of no allelic imbalance (i.e. with *a* fixed to *a* = 0) to compute p-values. We correct the p-values for multiple testing using the Benjamini-Hochberg method. To make it clear when we are referring to allelic imbalance in RNA instead of DNA, we refer to *a* for RNA-seq reads as *a*_RNA_.

### Gene ontology enrichment analysis

Gene ontology (GO) enrichment analysis for Biological Processes (BP) was performed using topGO (2.34.0). Enrichment was calculated using Fisher’s Exact Test and all genes tested for ASE in neuroblastoma were used as the universe.

### Correlation between DNA allele-imbalance and SNP-array predictions

We used GenomicRanges (1.34.0) to find overlaps between SCNAs detected using our DNA allelic-imbalance method and SCNA predictions obtained from TARGET which were generated using HumanHap 550K Beadchip (SNP-array)^39^. For segments which show at least a 50% overlap, we computed Spearman’s correlation between segmented DNA allele imbalance i.e., δ_CBS_ and corrected Log R ratio estimated using SNP-array data from TARGET^39^.

### Association between allele-specific expression and SCNAs

We assigned our candidate genes to genomic segments predicted to be SCNAs based on the location of their promoters (transcription start site +/- 1500bp) using GenomicRanges (1.34.0)^68^ and computed Spearman’s correlation between ASE (a_RNA_) and SCNA score. We corrected the p-values for multiple testing using the Benjamini-Hochberg procedure. We only tested genes with non-zero variance in both SCNA scores and a_RNA_, and with an SCNA score in at least one sample ≥ 0.09 . Karyotype plots for Spearman’s correlation coefficient were generated using ggbio (1.30.0)^69^.

### Correlations between allele-specific expression and promoter methylation

In Human Methylation 450K BeadChIP array (HM450K) data the ratios of intensities between methylated and unmethylated CpG probes are referred to as Beta values (β) and range from 0 (unmethylated) to 1 (completely methylated). We downloaded a pre-computed β matrix for 87 neuroblastoma samples from TARGET and annotated the CpG probe positions based on GENCODE (version 28) genes. Then we computed the mean β for promoter regions (transcription start site +/- 1500 bp) and computed Spearman’s correlations between promoter methylation and ASE (a_RNA_) for 1,049 NB-ASE genes. We corrected the p-values for multiple testing using the Benjamini Hochberg procedure. Under an FDR of 0.1 we found 6 genes with a significant association between DNA methylation and ASE (Suppl table 7). We only tested genes with at least 3 CpG probes within promoter regions.

### Survival and expression analysis in neuroblastoma patients

We analyzed the SEQC/MAQC-III Consortium dataset consisting of 498 individuals using the R2: Genomics Analysis and Visualization Platform (http://r2.amc.nl) to generate Kaplan-Meier survival plots for neuroblastoma^52^. We also downloaded normalized gene expression matrix (i.e. log (1 + FPKM)) for SEQC/MAQC-III Consortium dataset from Gene Expression Omnibus (GSE49711)^52^ and generated gene expression heatmaps using ComplexHeatmap (1.20.0) and ggplot2 (3.2.1)^67, 70^.

### Cell culture and transfection

The SK-N-BE(2) and SK-N-SH cell lines were purchased from the American Type Culture Collection (www.atcc.org) and grown in a humidified chamber with 5% CO2 in RPMI 1640 medium (Gibco, #11875119) supplemented with 10% fetal bovine serum (FBS), 2mM L-glutamine, sodium pyruvate, non-essential amino acids and 1% antibiotic antimycotic.

For stable transfection, the cell lines were seeded in 6-well plates and allowed to grow to 70% confluence. Cells were transfected with shRNAs purchased from Sigma-Aldrich (shPTPRH-373, #TRCN0000355579; shPTPRH-1136, #TRCN0000355581; shPTPRH-1947, #TRCN0000355580; shPTPRH-3265, TRCN0000355631; shPTPRH-3621, #TRCN0000002866) with jetOPTIMUS® DNA transfection Reagent (VWR, #76299-632) following the protocol provided by the manufacturer. The cells were selected in RPMI containing puromycin 24hrs after transfection. (SK-N-BE(2), 1μg/ml puromycin and SK-N-SH, 1.25μg/ml puromycin). The stably transfected cells were maintained in complete medium supplemented with puromycin until use.

### Quantitative RT-PCR analysis

SK-N-BE(2) and SK-N-SH cells were seeded in 6-well plates and allowed to grow to 50% confluence. RNA was isolated from the cells with Zymo Quick-RNA Miniprep kit (VWR, #76299-632) according to the manufacturer’s instructions. Total RNA was reversed transcribed into cDNA using High-Capacity cDNA Reverse Transcription Kit (Fisher Scientific, #4374966). Real-time PCR was performed using iTaq™ Universal SYBR Green Supermix (Bio-Rad Laboratories, #1725122) on a Bio-Rad CFX96 system. Gene expression was analyzed by the log2ΔΔCt method.

### Immunoblotting

SK-N-BE(2) and SK-N-SH cells were seeded in 6-well plates and allowed to grow to 70-80% confluence. Cells were lysed in RIPA buffer, and the protein concentration was determined by a Pierce BCA protein assay (Life Technologies, #23225). Equal amounts of protein were loaded into 8% Bolt™ Bis-Tris Plus gels (Life Technologies, #NW00085BOX), separated by SDS-PAGE and then transferred to PVDF membranes. The membranes were incubated with primary antibodies (PTPRH antibody, 1:1000, Fisher Scientific, #PIPA531340) overnight at 4°C. The membranes were then probed with appropriate horseradish peroxidase-conjugated secondary antibodies (Goat Anti-Rabbit IgG(H+L)-HRP Conjugate, Bio-rad Laboratories, #170- 6515). The immunoblots were visualized with SuperSignal West Pico Plus Chemiluminescent Substrate (Life Technologies, #PI34580).

### Proliferation and migration assays

SK-N-BE(2) and SK-N-SH cells were plated in 96-well plates at a seeding density of 7,500 cells/well and allowed to attach overnight. Cells were then monitored by continuous live-cell imaging in the IncuCyte® Zoom^TM^ system (Essen Bioscience) and 10x phase contrast images were taken every 3hrs. Cell confluence in each image was calculated by the IncuCyte® analysis software.

For migration assays, SK-N-BE(2) and SK-N-SH cells were seeded in IncuCyte® Imagelock 96-well plates (Essen Bioscience, #4379) at seeding densities between 100,000-200,000 cells/well and allowed to grow to 100% confluence. Cells were treated with 10μM Cytosine arabinoside (Sigma-Aldrich, #C1768) for 4hrs, and then identical scratch wounds were made in each well with a 96-pin WoundMaker (Essen Bioscience). Wound closure was monitored by continuous live-cell imaging in the IncuCyte Zoom^TM^ (Essen Bioscience) and 10x phase contrast images were taken every 6hrs. The wound closure percentage was calculated using IncuCyte Scratch Wound Analysis Software.

To analyze the cellular proliferation and migration rates, we computed the mean confluence and mean wound closure at each time point across replicates. To allow for non-linearity in both types of data, we performed cubic spline regression with 3 knots (two degrees of freedom) to describe the change in confluence or wound closure with time. To quantify differences in proliferation or migration between knockdown and parental cells, we included an interaction term between knockdown status and the time spline in the model. Specifically, we utilized the following linear model command in R, where ‘response’ is the confluence or wound closure, ‘ns’ is the cubic spline function, and ‘cond’ is a factor giving knockdown status (parental or knockdown):

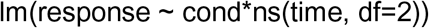

Significance of the interaction term was determined with an F-test.

## Acknowledgements

The results published here are in whole or part based upon data generated by the Therapeutically Applicable Research to Generate Effective Treatments (TARGET) initiative, phs000218, managed by the NCI. The data used for this analysis are available (phs000467.v1.p1). Information about TARGET can be found at http://ocg.cancer.gov/programs/target.

The Genotype-Tissue Expression (GTEx) Project was supported by the Common Fund of the Office of the Director of the National Institutes of Health, and by NCI, NHGRI, NHLBI, NIDA, NIMH, and NINDS. The data used for the analyses described in this manuscript were obtained from dbGaP accession number phs000424.v7.p2 on 07/16/2018.

This research was supported by the National Cancer Institute funded Salk Institute Cancer Center (NIH/NCI CCSG: P30 014195), and a grant from Padres Pedal the Cause/RADY #PTC2019 awarded to G.M. and J.E.

A.S. was supported by a Pioneer Fund Postdoctoral Scholar Award. G.M. was supported by the Frederick B. Rentschler Developmental Chair.

## Author contributions

G.M., J.E. and A.S. conceived of the project. G.M. and J.E. obtained funding for the project. Data processing and analysis was performed by A.S. A.S. wrote the initial draft and A.S. and G.M. wrote the manuscript. P.Z. provided comments on the manuscript. G.M. supervised the analysis and data processing. Y.H. performed all experiments under the supervision of P.Z.

## Supplementary Figures

**Supplementary Figure S1:**
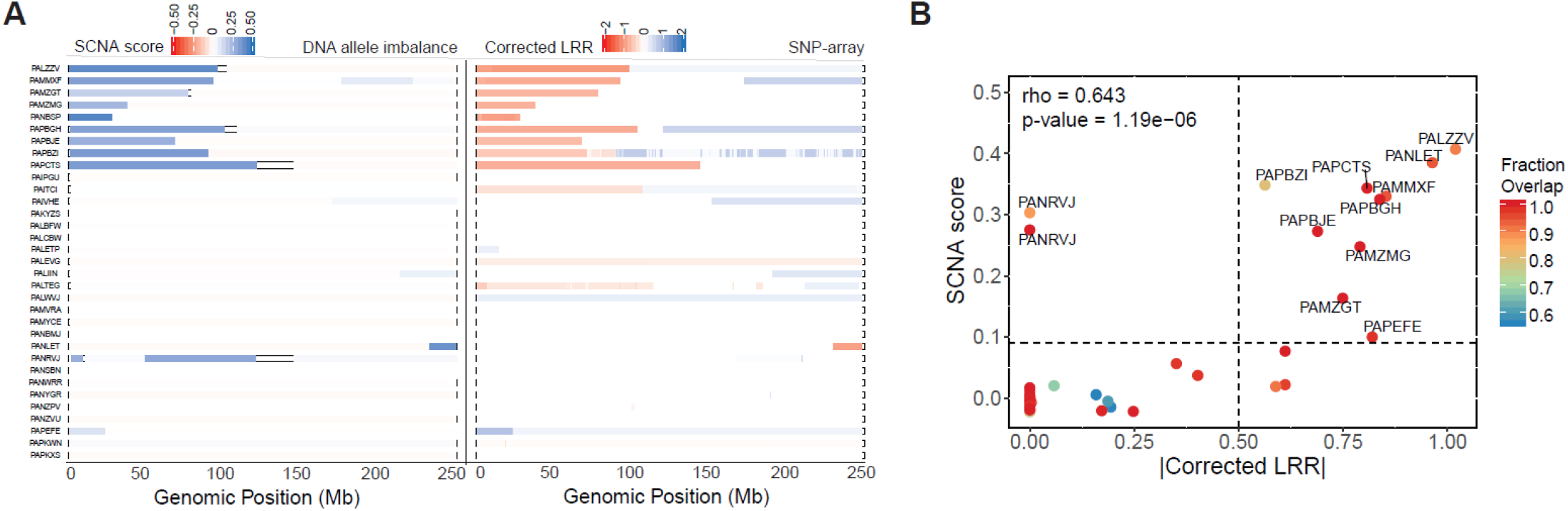
Validation of SCNA scores using SNP-array data. **A)** SCNA predictions based on DNA allelic imbalance compared to SNP-array predictions. Left panel: SCNA scores across chromosome 1 estimated from DNA allelic imbalance from 33 neuroblastoma patients. Right Panel: Corrected Log R ratio (or Corrected LRR) calculated from SNP array data for the same 33 patients. Corrected LRR is defined as aneuploidy corrected total probe intensity of a given genomic segment relative to a canonical set of normal controls. **B)** Spearman’s rank correlation between SCNA score and absolute Corrected LRR for chromosome 1 (Spearman’s correlation coefficient = 0.64, p-value = 1.19e-06). Exome-seq and SNP arrays may use slightly different set of SNPs to predict SCNAs. Therefore, the Circular Binary Segmentation (CBS) algorithm tends to output segments which do not share the same genomic start and end positions. To be able to directly compare SCNA detection using DNA allele imbalance and SNP array, we first calculated the fraction overlap between genome segments identified by the respective methods. Next, we performed pairwise Spearman’s correlation between SCNA score and absolute Corrected LRR for genomic segments with fraction overlap ≥ 0.5 or 50%. The points in the correlation scatter plot are colored by fraction overlap. The two points labelled PANRVJ correspond to two disjointed SCNAs spanning chr1:1922327-9171333 and chr1:49201909-120298048. These regions showed absolute Corrected LRR < 0.5 and were annotated as copy neutral by SNP array. We suspect that these segments may be copy-neutral loss of heterozygosity regions, which are not detectable using direct analysis of SNP-array intensities.

**Supplementary Figure S2:**
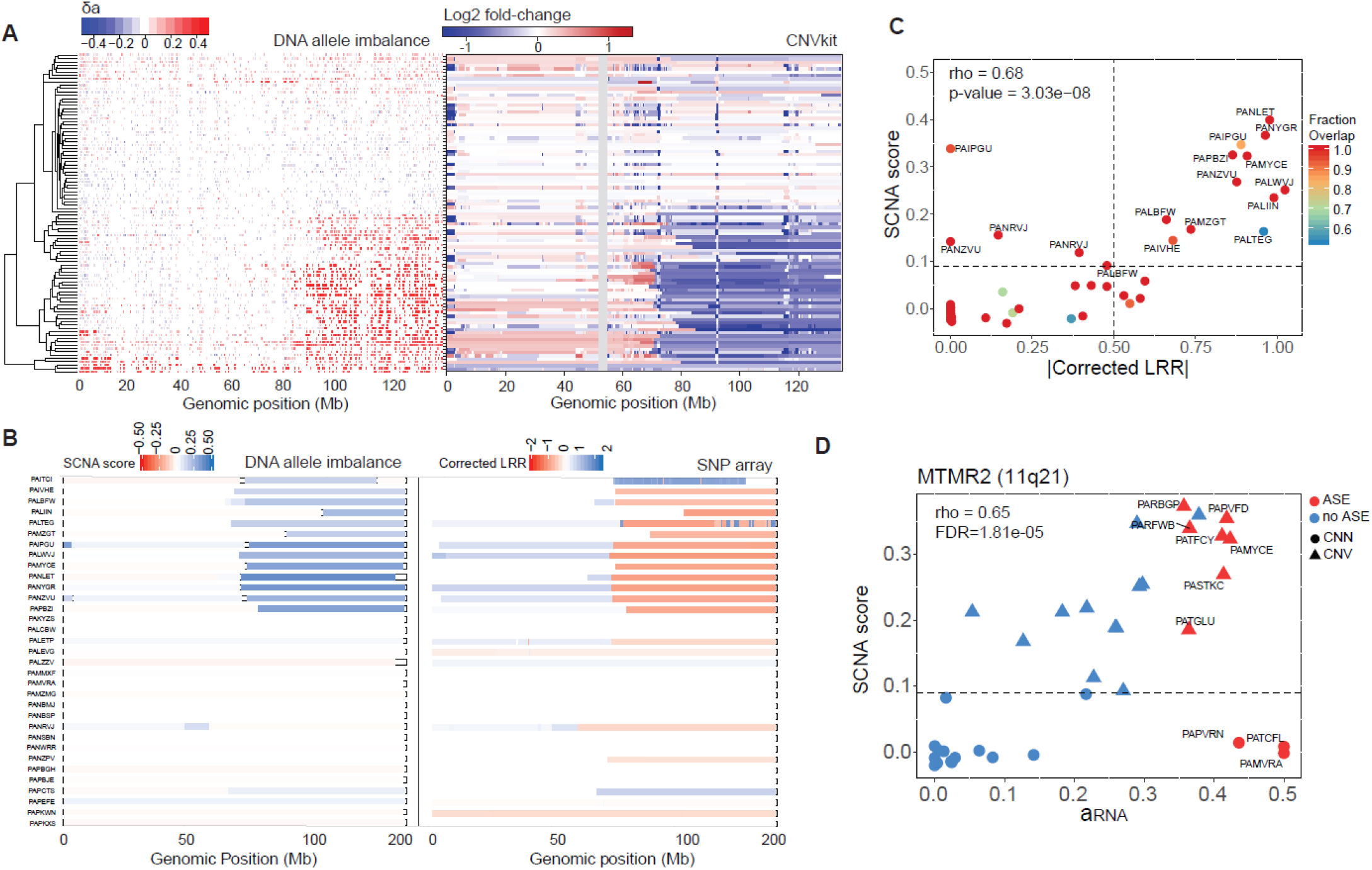
SCNA predictions for chromosome 11. **A**) Comparison between DNA- imbalance SCNA predictions and CNVkit predictions for chr11. Panel 1: Heatmap of difference in DNA allele-imbalance between tumor and normal tissue (i.e., δ_a_) for 96 neuroblastoma patients. Panel 2: Log2 fold-change in normalized read coverage between tumor and normal tissues estimated using CNVkit. **B**) Comparison between DNA-imbalance predictions and SNP-array predictions for chr11. Panel 1: SCNA score across 33 neuroblastoma patients with SNP-array data in TARGET. Panel 2: corrected LRR calculated array SNP-array available through TARGET. **C**) Spearman’s rank correlation between SCNA score and absolute corrected LRR for chr11 (Spearman’s correlation coefficient = 0.68, p-value = 3.03e-08). The points in the correlation scatter plot are colored by fraction overlap. **D**) Spearman’s rank correlation between ASE (a_RNA_) and SCNA score for *MTMR2*, a gene located within a common deletion segment on cytoband 11q21 (Spearman’s correlation coefficient = 0.65, p-value= 1.81e-05).

**Supplementary Figure S3:**
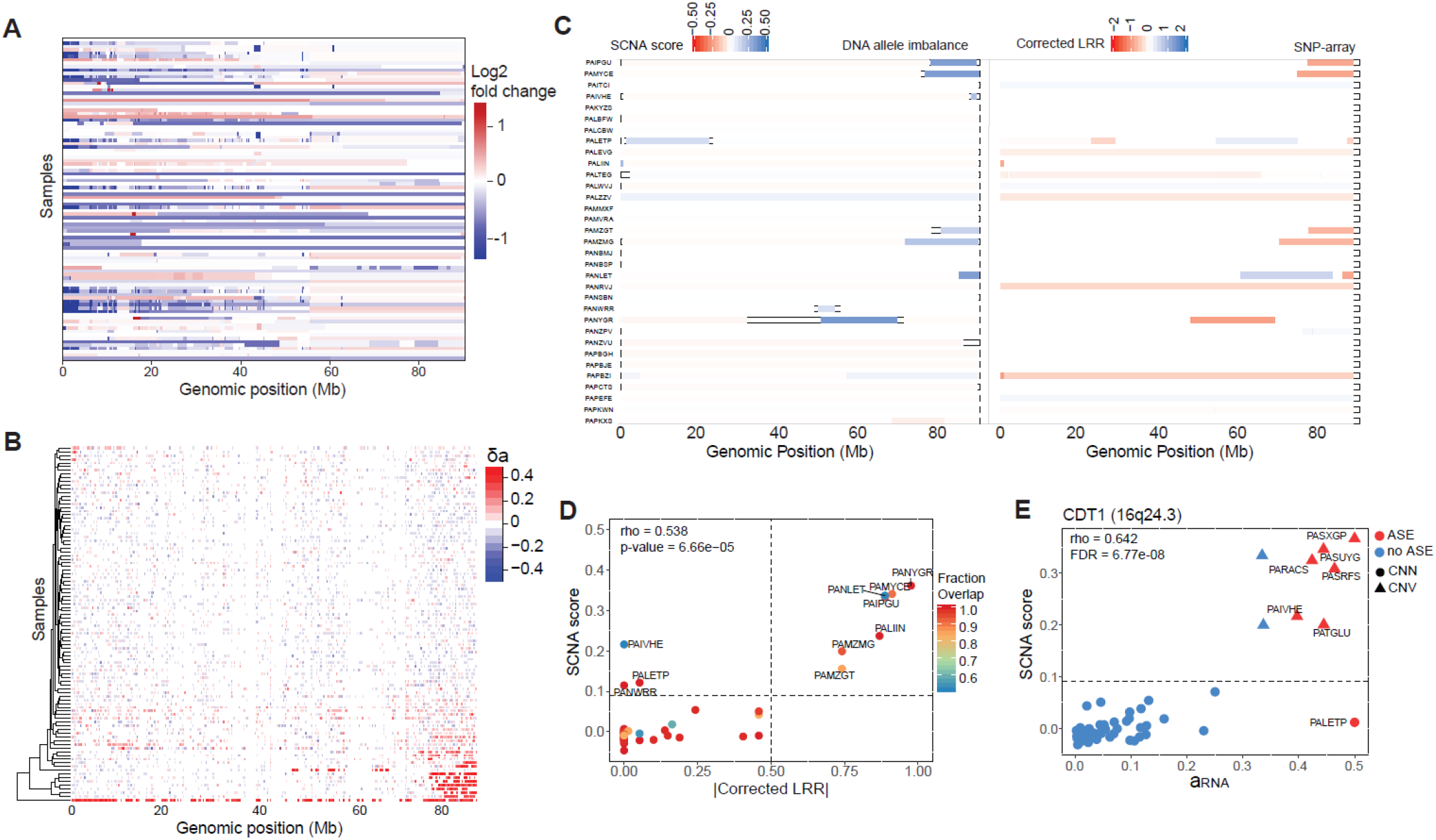
Detection of SCNAs on chromosome 16. **A**) Log2 fold-change in normalized read coverage between tumor and normal tissues estimated using CNVkit for chromosome 16. These predictions are noisy and it difficult to separate true signal from background. **B**) Heatmap of δ_a_ for chromosome 16. Chromosome 16q SCNAs are observed in 18 samples and 16p SCNAs are observed in 4 samples. **C**) Comparison between DNA-imbalance and SNP-array predictions for chromosome 16 for 33 neuroblastoma samples indicates good concordance between our predictions and SNP-array predictions. **D**) Spearman’s rank correlation between SCNA score and absolute corrected LRR for chromosome 16 (Spearman’s correlation coefficient = 0.54, p-value = 6.7e-05). The points in the correlation scatter plot are colored by fraction overlap. **E**) Spearman’s rank correlation between ASE (a_RNA_) and SCNA score for *CDT1*, a gene located in distal region of the q-arm (i.e. 16q24.3) (Spearman’s correlation coefficient = 0.64, FDR corrected p-value= 6.77e- 08).

**Supplementary Figure S4:**
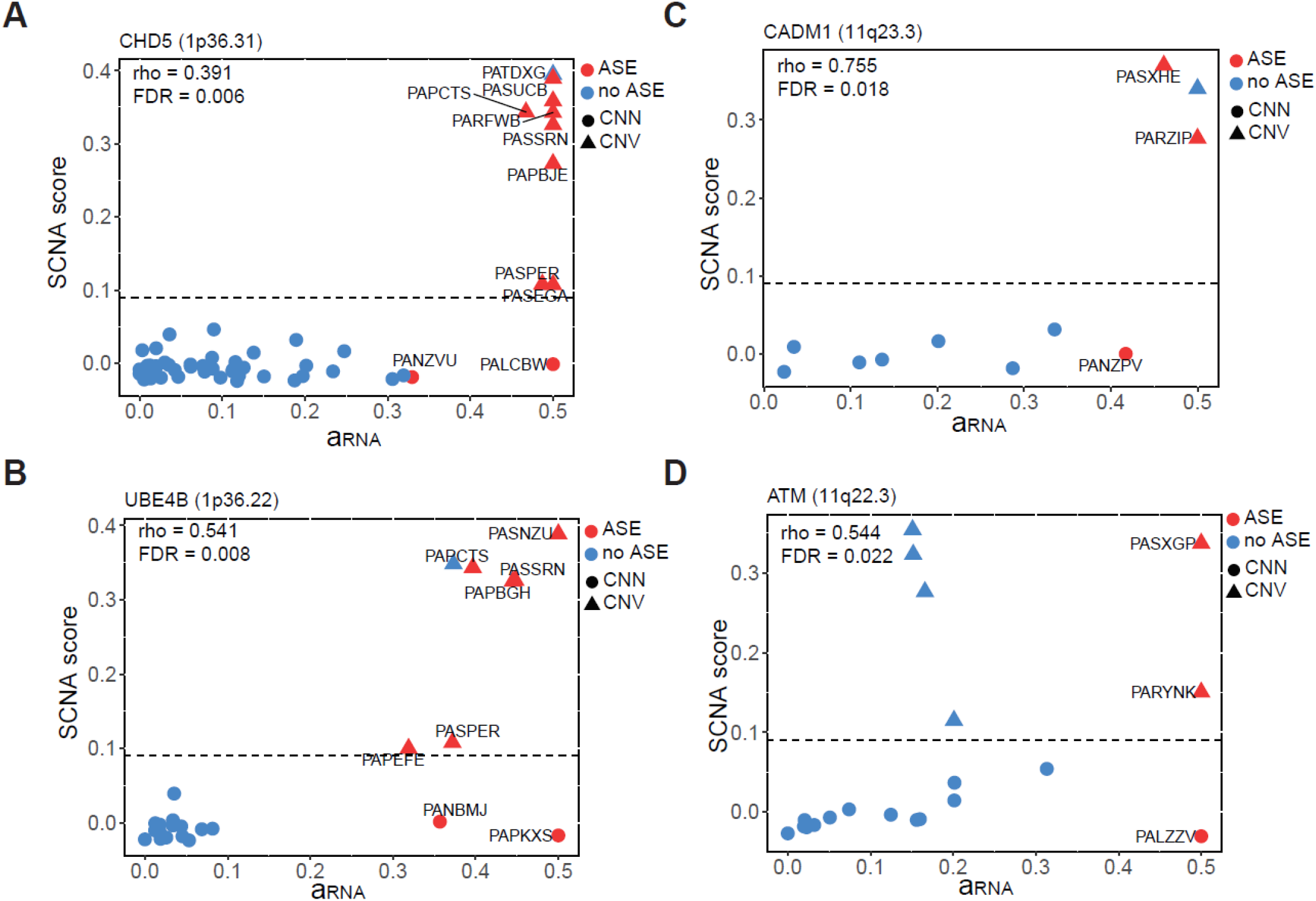
Haplo-insufficient tumor suppressors within common SCNAs may be dysregulated by secondary mechanisms. **A**-**C**) Spearman’s correlation between ASE (a_RNA_) and SCNA score for 1p deletion genes *CHD5* (**A**) and *UBE4B* (**B**), and 11q deletion genes *CADM1* (**C**) and *ATM* (**D**). Several samples show ASE in the absence of SCNAs.

**Supplementary Figure S5:**
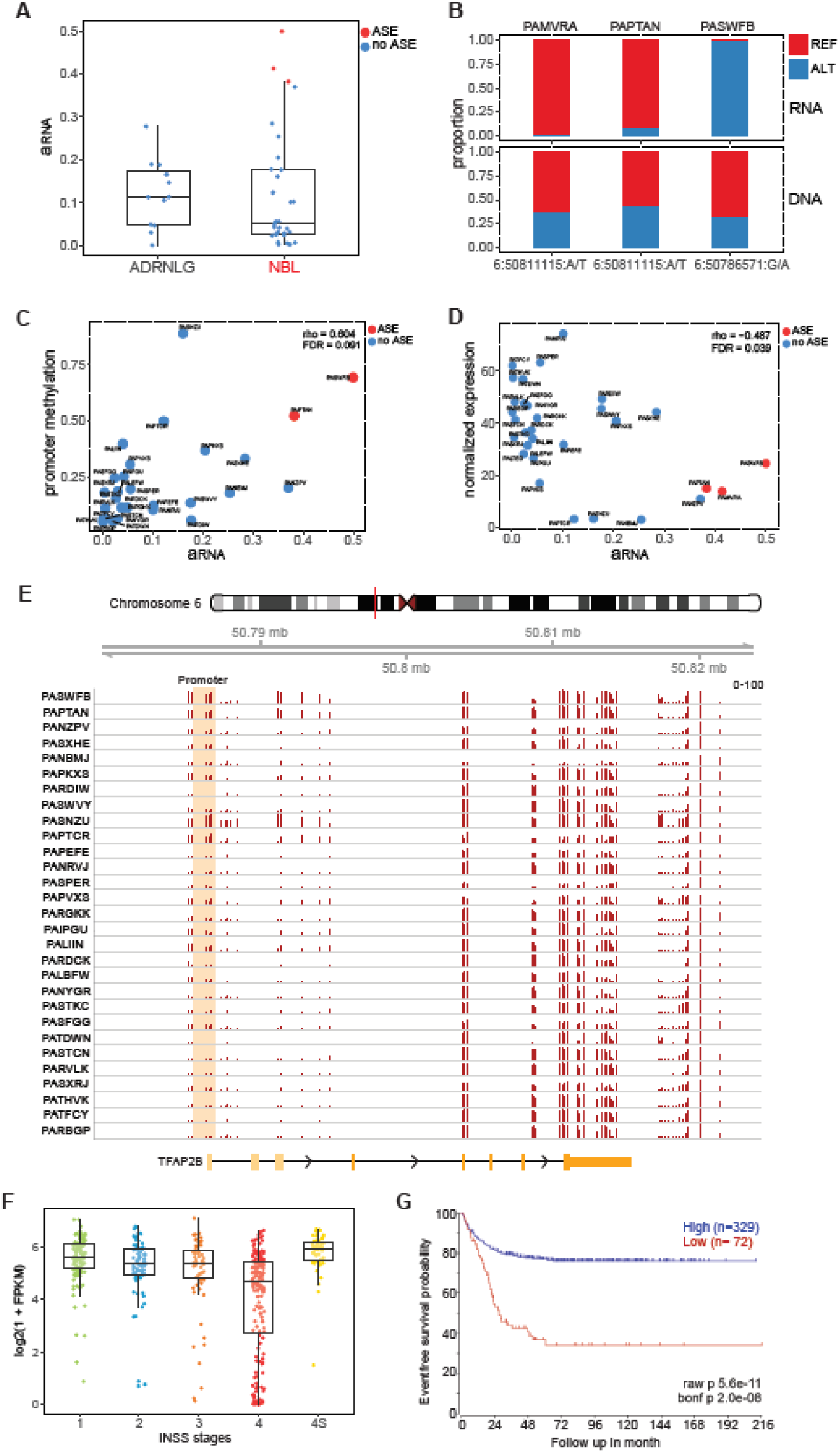
*TFAP2B* expression in neuroblastoma is regulated by promoter methylation. **A**) ASE (a_RNA_) of *TFAP2B* in neuroblastoma and adrenal gland tissues. **B**) Reference and alternate allele proportion for RNA-seq and exome-seq reads at heterozygous sites which were used to estimate ASE for *TFAP2B*. **C**) Spearman’s correlation between ASE (a_RNA_) and gene expression for *TFAP2B.* **D**) Spearman’s correlation between ASE (a_RNA_) and promoter methylation for *TFAP2B*. DNA methylation data was missing for 1 neuroblastoma sample (PAMVRA). **E**) Genomic distribution of HM450K β-values for *TFAP2B* locus. The *TFAP2B* promoter is highlighted (yellow box). **F**) Expression profile of *TFAP2B* across different stages of disease for 498 neuroblastoma patients obtained from SEQC/MAQC-III Consortium data set. We observed loss of expression of *TFAP2B* in stage 4 or metastatic disease suggesting this gene might act as a tumor suppressor. **G**) Kaplan Meier survival analysis for MYCN non-amplified patients from the SEQC/MAQC-III Consortium data set.

**Supplementary Figure S6:**
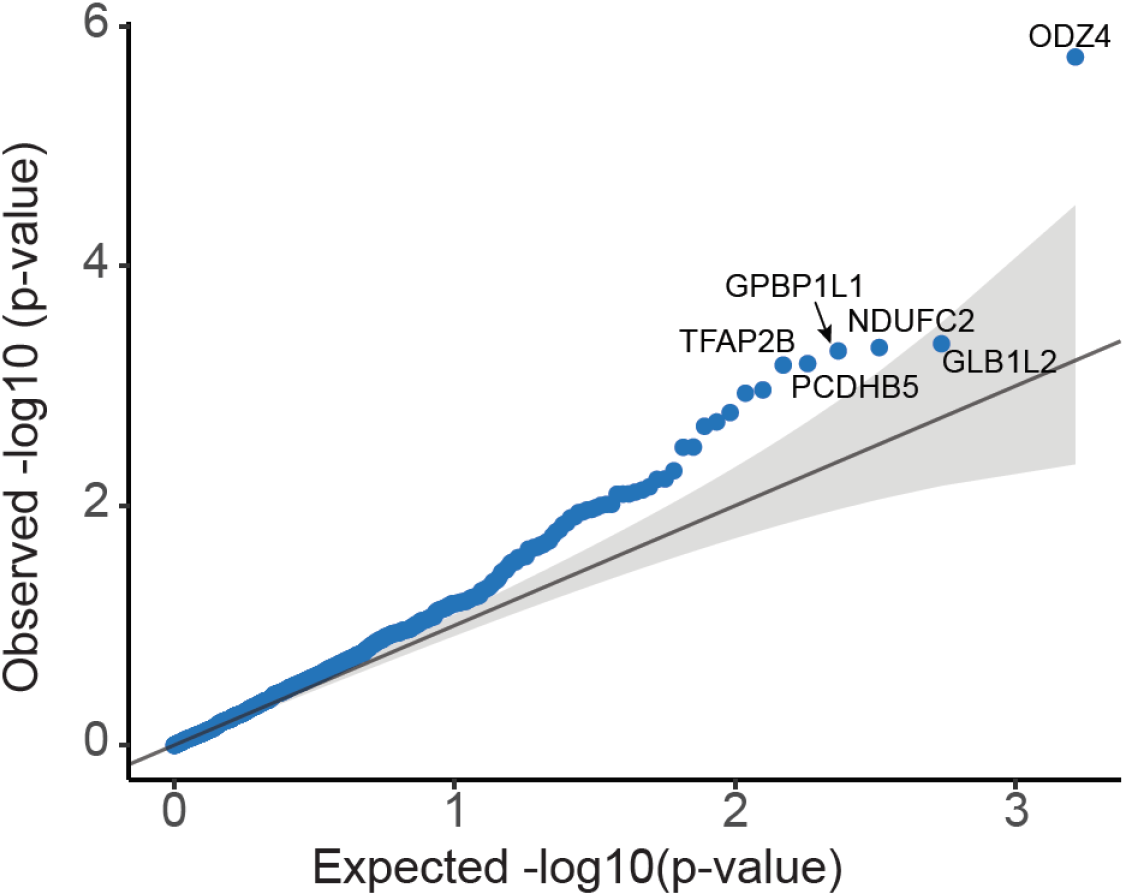
Quantile-quantile plot for Spearman’s correlation analysis between ASE (a_RNA_) and promoter methylation for 1,049 NB-ASE genes. Under an FDR of 10% only 6 genes were significantly correlated.

**Supplementary Figure S7:**
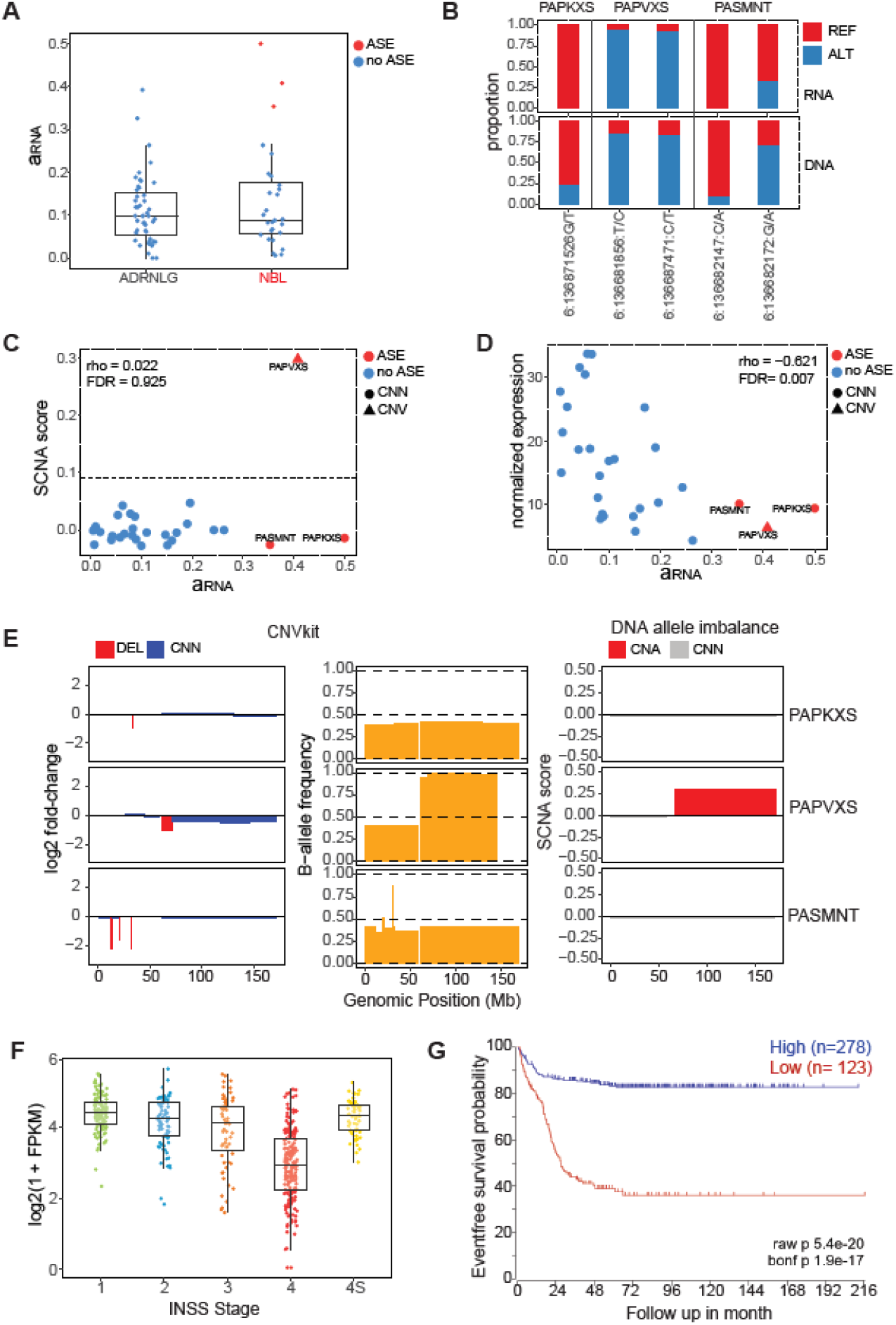
Regulation of *MAP7* gene-expression in neuroblastoma. **A)** ASE (a_RNA_) of *MAP7* in neuroblastoma and adrenal gland. **B)** Reference and alternate allele proportion for RNA-seq and exome-seq reads at heterozygous sites which were used to estimate ASE for *MAP7*. **C)** Spearman’s correlation between ASE (a_RNA_) and SCNA score for *MAP7*. **D)** Spearman’s correlation between absolute a_RNA_ and gene expression of *MAP7.* **E)** Comparison between our DNA allelic-imbalance predictions and CNVkit predictions for chromosome 6 for 3 neuroblastoma patients which showed significant ASE of *MAP7* (FDR corrected p-value ≤ 10%). Panel 1 shows the log2 fold-change in normalized coverage between tumor and normal samples estimated using CNVKit. Panel 2 shows normalized B-allele-frequencies (BAF) also calculated using CNVkit. Panel 3 shows SCNA scores measured using our method. Results from CNVKit and our method confirm the presence of a 6q SCNA in sample PAPVXS. **F)** Boxplot of normalized gene-expression of *MAP7* across different disease stages for 498 neuroblastoma patients from the SEQC/MAQC-III Consortium data. **G)** Kaplan Meier survival analysis for MYCN non-amplified patients from the SEQC/MAQC-III Consortium data set.

**Supplemental Figure S8:**
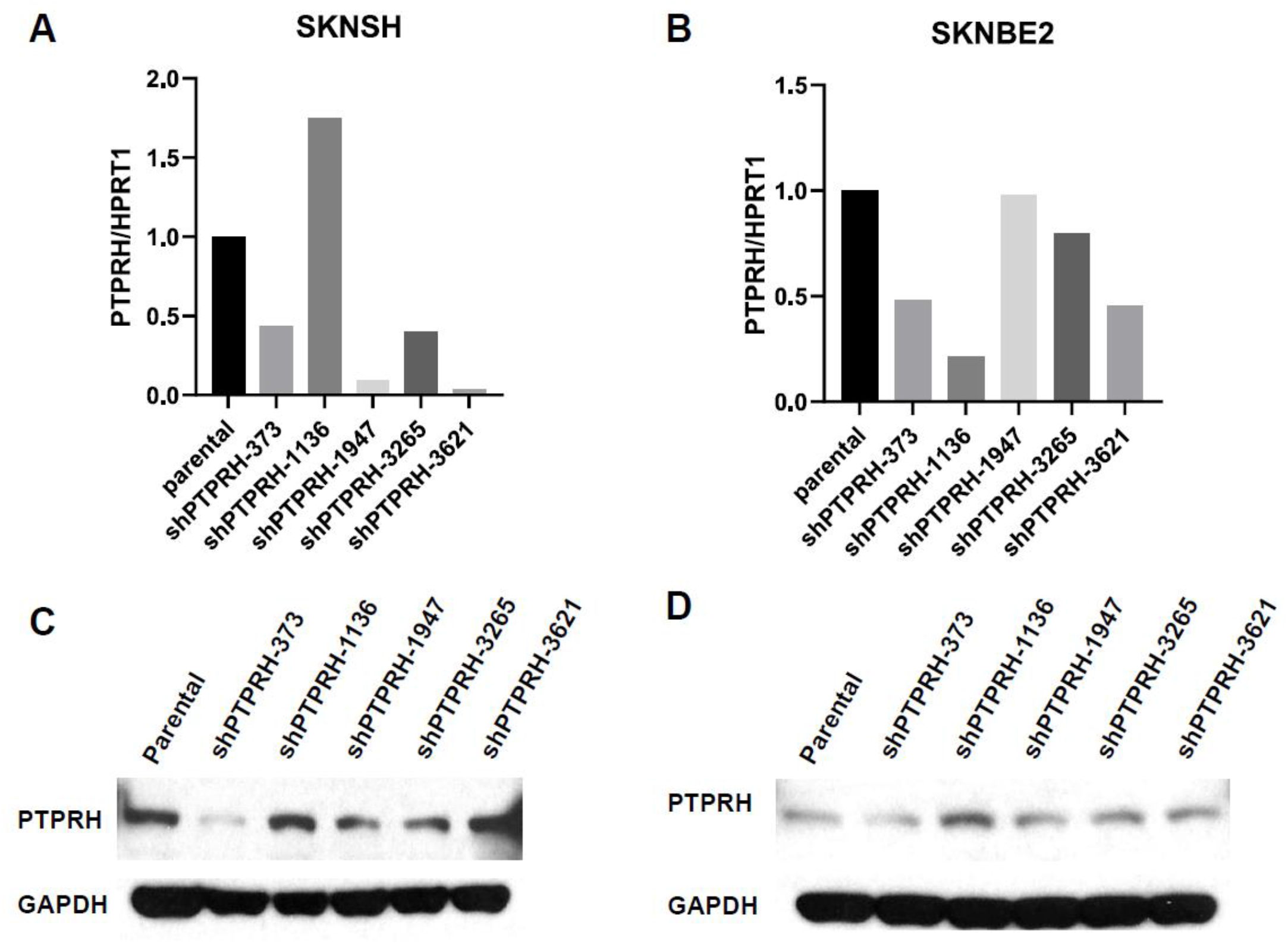
Knockdown of PTPRH in neuroblastoma cell lines. A-B) qPCR to measure *PTPRH* expression in SK-N-SH (**A**) or SK-N-BE(2) (**B**) neuroblastoma cell lines stably transfected with 5 different shRNAs targeting *PTPRH*. Gene expression is plotted as 2^-ΔΔCt^ normalized to *HPRT1* expression. **C**-Western blot of *PTPRH* and *GAPDH* protein expression for the same SK-N-SH (**C**) and SK-N-BE(2) (**D**) cell lines. shRNA shPTPRH-373 consistently reduced gene and protein expression of *PTPRH* in both SK-N-SH and SK-N-BE(2) cells and was used for all downstream experiments.

## Supplementary tables

**Supplementary table 1:** Results from Allele-Specific Expression (ASE) analysis of RNA-seq data for 96 neuroblastoma samples from TARGET.

**Supplementary table 2:** Number of significant (FDR ≤ 0.1 or 10%) and testable (i.e. at least 1 heterozygous site with ≥ 10 reads) samples for neuroblastoma tumors, adrenal gland, and whole-blood tissues for all significant ASE genes. Genes were considered to have neuroblastoma-specific ASE if they met these criteria; a) testable in ≥ 10 neuroblastoma and normal (i.e., adrenal-gland and whole-blood) samples and b) significant in ≥3 neuroblastoma and significant ≤ 1 normal sample.

**Supplementary table 3:** Top 20 Gene Ontology (Biological processes) categories enriched for 1,049 NB-ASE genes.

**Supplementary table 4:** SCNA scores for 96 neuroblastoma patient samples.

**Supplementary table 5:** Spearman’s correlation between ASE (a_RNA_) for 1,049 NB-ASE genes and SCNA score for overlapping genomic segment.

**Supplementary table 6:** Annotated table of high-impact somatic mutations detected using exome-seq data in 96 neuroblastoma tumor-normal pairs.

**Supplementary table 7:** Spearman correlation between ASE (a_RNA_) and gene expression (z-score normalized FPKM) for 1,049 NB-ASE genes.

**Supplementary table 8:** Spearman correlation between ASE (a_RNA_) and mean promoter methylation for 1,049 NB-ASE genes.

## References

1. Mlakar, V. et al. 11q deletion in neuroblastoma: a review of biological and clinical implications. Mol Cancer 16, 114 (2017).

2. Maris, J.M., Hogarty, M.D., Bagatell, R. & Cohn, S.L. Neuroblastoma. Lancet 369, 2106–20 (2007).

3. Schleiermacher, G., Janoueix-Lerosey, I. & Delattre, O. Recent insights into the biology of neuroblastoma. Int J Cancer 135, 2249–61 (2014).

4. Bagatell, R. & Cohn, S.L. Genetic discoveries and treatment advances in neuroblastoma. Curr Opin Pediatr 28, 19–25 (2016).

5. Pinto, N.R. et al. Advances in Risk Classification and Treatment Strategies for Neuroblastoma. J Clin Oncol 33, 3008–17 (2015).

6. Matthay, K.K., et al. Neuroblastoma. Nat Rev Dis Primers 2, 16078 (2016).

7. Pugh, T.J. et al. The genetic landscape of high-risk neuroblastoma. Nat Genet 45, 279–84 (2013).

8. Cheung, N.K. & Dyer, M.A. Neuroblastoma: developmental biology, cancer genomics and immunotherapy. Nat Rev Cancer 13, 397–411 (2013).

9. Huang, M. & Weiss, W.A. Neuroblastoma and MYCN. Cold Spring Harb Perspect Med 3, a014415 (2013).

10. Maris, J.M. et al. Loss of heterozygosity at 1p36 independently predicts for disease progression but not decreased overall survival probability in neuroblastoma patients: a Children’s Cancer Group study. J Clin Oncol 18, 1888–99 (2000).

11. Plantaz, D. et al. Gain of chromosome 17 is the most frequent abnormality detected in neuroblastoma by comparative genomic hybridization. Am J Pathol 150, 81–9 (1997).

12. Attiyeh, E.F. et al. Chromosome 1p and 11q deletions and outcome in neuroblastoma. N Engl J Med 353, 2243–53 (2005).

13. Zage, P.E. et al. UBE4B levels are correlated with clinical outcomes in neuroblastoma patients and with altered neuroblastoma cell proliferation and sensitivity to epidermal growth factor receptor inhibitors. Cancer 119, 915–23 (2013).

14. Garcia, I. et al. Expression of the neuron-specific protein CHD5 is an independent marker of outcome in neuroblastoma. Mol Cancer 9, 277 (2010).

15. Kolla, V. et al. The tumour suppressor CHD5 forms a NuRD-type chromatin remodelling complex. Biochem J 468, 345–52 (2015).

16. Yang, H.W. et al. Genomic structure and mutational analysis of the human KIF1B gene which is homozygously deleted in neuroblastoma at chromosome 1p36.2. Oncogene 20, 5075–83 (2001).

17. Liu, Z. et al. CASZ1, a candidate tumor-suppressor gene, suppresses neuroblastoma tumor growth through reprogramming gene expression. Cell Death Differ 18, 1174–83 (2011).

18. Schlisio, S. et al. The kinesin KIF1Bbeta acts downstream from EglN3 to induce apoptosis and is a potential 1p36 tumor suppressor. Genes Dev 22, 884–93 (2008).

19. Fujita, T. et al. CHD5, a tumor suppressor gene deleted from 1p36.31 in neuroblastomas. J Natl Cancer Inst 100, 940–9 (2008).

20. Welch, C., Chen, Y. & Stallings, R.L. MicroRNA-34a functions as a potential tumor suppressor by inducing apoptosis in neuroblastoma cells. Oncogene 26, 5017–22 (2007).

21. Henrich, K.O. et al. CAMTA1, a 1p36 tumor suppressor candidate, inhibits growth and activates differentiation programs in neuroblastoma cells. Cancer Res 71, 3142–51 (2011).

22. Mohammadi, P. et al. Genetic regulatory variation in populations informs transcriptome analysis in rare disease. Science 366, 351–356 (2019).

23. Liu, Y. et al. Discovery of regulatory noncoding variants in individual cancer genomes by using cis-X. Nat Genet 52, 811–818 (2020).

24. Pastinen, T. Genome-wide allele-specific analysis: insights into regulatory variation. Nat Rev Genet 11, 533–8 (2010).

25. Skelly, D.A., Johansson, M., Madeoy, J., Wakefield, J. & Akey, J.M. A powerful and flexible statistical framework for testing hypotheses of allele-specific gene expression from RNA-seq data. Genome Res 21, 1728–37 (2011).

26. Knowles, D.A. et al. Allele-specific expression reveals interactions between genetic variation and environment. Nat Methods 14, 699–702 (2017).

27. Buckberry, S., Bianco-Miotto, T., Hiendleder, S. & Roberts, C.T. Quantitative allele-specific expression and DNA methylation analysis of H19, IGF2 and IGF2R in the human placenta across gestation reveals H19 imprinting plasticity. PLoS One 7, e51210 (2012).

28. Deng, Q., Ramskold, D., Reinius, B. & Sandberg, R. Single-cell RNA-seq reveals dynamic, random monoallelic gene expression in mammalian cells. Science 343, 193–6 (2014).

29. Chess, A. Mechanisms and consequences of widespread random monoallelic expression. Nat Rev Genet 13, 421–8 (2012).

30. Krueger, C. & Morison, I.M. Random monoallelic expression: making a choice. Trends Genet 24, 257–9 (2008).

31. Rachmilewitz, J. et al. Parental imprinting of the human H19 gene. FEBS Lett 309, 25–8 (1992).

32. Munirajan, A.K. et al. KIF1Bbeta functions as a haploinsufficient tumor suppressor gene mapped to chromosome 1p36.2 by inducing apoptotic cell death. J Biol Chem 283, 24426–34 (2008).

33. Mayba, O. et al. MBASED: allele-specific expression detection in cancer tissues and cell lines. Genome Biol 15, 405 (2014).

34. Van Loo, P. et al. Allele-specific copy number analysis of tumors. Proc Natl Acad Sci U S A 107, 16910–5 (2010).

35. Yao, R. et al. Evaluation of three read-depth based CNV detection tools using whole-exome sequencing data. Mol Cytogenet 10, 30 (2017).

36. Zare, F., Dow, M., Monteleone, N., Hosny, A. & Nabavi, S. An evaluation of copy number variation detection tools for cancer using whole exome sequencing data. BMC Bioinformatics 18, 286 (2017).

37. Olshen, A.B., Venkatraman, E.S., Lucito, R. & Wigler, M. Circular binary segmentation for the analysis of array-based DNA copy number data. Biostatistics 5, 557–72 (2004).

38. Talevich, E., Shain, A.H., Botton, T. & Bastian, B.C. CNVkit: Genome-Wide Copy Number Detection and Visualization from Targeted DNA Sequencing. PLoS Comput Biol 12, e1004873 (2016).

39. Attiyeh, E.F. et al. Genomic copy number determination in cancer cells from single nucleotide polymorphism microarrays based on quantitative genotyping corrected for aneuploidy. Genome Res 19, 276–83 (2009).

40. Grundy, P.E. et al. Loss of heterozygosity for chromosomes 1p and 16q is an adverse prognostic factor in favorable-histology Wilms tumor: a report from the National Wilms Tumor Study Group. J Clin Oncol 23, 7312–21 (2005).

41. Uryu, K. et al. Identification of the genetic and clinical characteristics of neuroblastomas using genome-wide analysis. Oncotarget 8, 107513–107529 (2017).

42. Altura, R.A. et al. Novel regions of chromosomal loss in familial neuroblastoma by comparative genomic hybridization. Genes Chromosomes Cancer 19, 176–84 (1997).

43. Koyama, H. et al. Mechanisms of CHD5 Inactivation in neuroblastomas. Clin Cancer Res 18, 1588–97 (2012).

44. Michels, E. et al. CADM1 is a strong neuroblastoma candidate gene that maps within a 3.72 Mb critical region of loss on 11q23. BMC Cancer 8, 173 (2008).

45. Mandriota, S.J. et al. Ataxia-telangiectasia mutated (ATM) silencing promotes neuroblastoma progression through a MYCN independent mechanism. Oncotarget 6, 18558–76 (2015).

46. Koldobskiy, M.A. et al. p53-mediated apoptosis requires inositol hexakisphosphate kinase-2. Proc Natl Acad Sci U S A 107, 20947–51 (2010).

47. Shames, D.S. & Minna, J.D. IP6K2 is a client for HSP90 and a target for cancer therapeutics development. Proc Natl Acad Sci U S A 105, 1389–90 (2008).

48. Popp, M.W. & Maquat, L.E. Nonsense-mediated mRNA Decay and Cancer. Curr Opin Genet Dev 48, 44–50 (2018).

49. Garcia-Lopez, J. et al. Large 1p36 Deletions Affecting Arid1a Locus Facilitate Mycn-Driven Oncogenesis in Neuroblastoma. Cell Rep 30, 454–464 e5 (2020).

50. Trochet, D. et al. Germline mutations of the paired-like homeobox 2B (PHOX2B) gene in neuroblastoma. Am J Hum Genet 74, 761–4 (2004).

51. Bourdeaut, F. et al. Germline mutations of the paired-like homeobox 2B (PHOX2B) gene in neuroblastoma. Cancer Lett 228, 51–8 (2005).

52. Zhang, W. et al. Comparison of RNA-seq and microarray-based models for clinical endpoint prediction. Genome Biol 16, 133 (2015).

53. Luscher, B., Mitchell, P.J., Williams, T. & Tjian, R. Regulation of transcription factor AP-2 by the morphogen retinoic acid and by second messengers. Genes Dev 3, 1507–17 (1989).

54. Ikram, F. et al. Transcription factor activating protein 2 beta (TFAP2B) mediates noradrenergic neuronal differentiation in neuroblastoma. Mol Oncol 10, 344–59 (2016).

55. Noguchi, T. et al. Inhibition of cell growth and spreading by stomach cancer-associated protein-tyrosine phosphatase-1 (SAP-1) through dephosphorylation of p130cas. J Biol Chem 276, 15216–24 (2001).

56. Takada, T. et al. Induction of apoptosis by stomach cancer-associated protein-tyrosine phosphatase-1. J Biol Chem 277, 34359–66 (2002).

57. Pirinen, M. et al. Assessing allele-specific expression across multiple tissues from RNA-seq read data. Bioinformatics 31, 2497–504 (2015).

58. Lee, S. et al. NGSCheckMate: software for validating sample identity in next-generation sequencing studies within and across data types. Nucleic Acids Res 45, e103 (2017).

59. Houtgast, E.J., Sima, V.M., Bertels, K. & Al-Ars, Z. Hardware acceleration of BWA-MEM genomic short read mapping for longer read lengths. Comput Biol Chem 75, 54–64 (2018).

60. McKenna, A. et al. The Genome Analysis Toolkit: a MapReduce framework for analyzing next-generation DNA sequencing data. Genome Res 20, 1297–303 (2010).

61. McLaren, W. et al. The Ensembl Variant Effect Predictor. Genome Biol 17, 122 (2016).

62. Dobin, A. et al. STAR: ultrafast universal RNA-seq aligner. Bioinformatics 29, 15–21 (2013).

63. Li, H. et al. The Sequence Alignment/Map format and SAMtools. Bioinformatics 25, 2078–9 (2009).

64. Liao, Y., Smyth, G.K. & Shi, W. featureCounts: an efficient general purpose program for assigning sequence reads to genomic features. Bioinformatics 30, 923–30 (2014).

65. Love, M.I., Huber, W. & Anders, S. Moderated estimation of fold change and dispersion for RNA-seq data with DESeq2. Genome Biol 15, 550 (2014).

66. van de Geijn, B., McVicker, G., Gilad, Y. & Pritchard, J.K. WASP: allele-specific software for robust molecular quantitative trait locus discovery. Nat Methods 12, 1061–3 (2015).

67. Gu, Z., Eils, R. & Schlesner, M. Complex heatmaps reveal patterns and correlations in multidimensional genomic data. Bioinformatics 32, 2847–9 (2016).

68. Lawrence, M. et al. Software for computing and annotating genomic ranges. PLoS Comput Biol 9, e1003118 (2013).

69. Yin, T., Cook, D. & Lawrence, M. ggbio: an R package for extending the grammar of graphics for genomic data. Genome Biol 13, R77 (2012).

70. SEQC/MAQC-III Consortium. A comprehensive assessment of RNA-seq accuracy, reproducibility and information content by the Sequencing Quality Control Consortium. Nat Biotechnol 32, 903–14 (2014).

